# Molecular mechanism of bacteriophage tail contraction-structure of an S-layer-penetrating bacteriophage

**DOI:** 10.1101/2023.08.04.551987

**Authors:** Jason S. Wilson, Louis-Charles Fortier, Robert P. Fagan, Per A. Bullough

**Author notes:** Present address: Biology Department, University of York, York, UK.

## Abstract

Viruses that infect bacteria (bacteriophages or phages) attach to the host cell envelope, inject their genetic material into the host cytosol and either persist as prophage or hijack the host machinery to produce progeny virions. Attachment is mediated through phage receptor binding proteins that are specific for different host cell surface molecules. A subset of phage, the myoviruses, possess contractile tails, the outer sheath of which contracts upon receptor binding, driving an inner tail tube through the cell envelope and delivering the phage genome into the host cytosol. The molecular details of phage tail contraction and mode of cell envelope penetration have remained poorly understood and were completely unknown for any phage infecting bacteria enveloped by a proteinaceous S-layer. Here we reveal the extended and contracted atomic structures of an intact contractile-tail phage that binds to and penetrates the protective S-layer of the Gram positive human pathogen *Clostridioides difficile*. Surprisingly, we find no evidence of the intrinsic enzymatic domains that other phages exploit in cell wall penetration, suggesting that sufficient energy is released upon tail contraction to penetrate the S-layer and the thick cell wall without enzymatic activity. However, it is also notable that the tail sheath subunits move less than those studied in related contractile injection systems such as the model phage T4. Instead, the unusually long tail length and flexibility upon contraction likely contribute towards the required free energy release for envelope penetration. Our results show that the principles of phage contraction and infection as determined in the model system of T4 are not universal. We anticipate that our structures will form a strong foundation to engineer *C. difficile* phages as therapeutics, and highlight important adaptations made in order to infect S-layer containing pathogens.

---

Phages are the most abundant biological entities on earth, yet only a tiny fraction have been characterised in any detail^1^. The most prevalent phage morphology described is that of an icosahedral capsid, containing the genome, attached to a tail^2^. The tail forms a conduit through which the genome is delivered into the host cell cytoplasm and is also associated with receptor binding functions^3^. Amongst phages, myoviruses have contractile tails of widely varying length, podoviruses have short non-contractile tails, while siphoviruses have long, flexible, non-contractile tails^4^.

Myovirus phage tails and bacterial tailocins both belong to the group of so-called contractile injection systems (CISs)^5^. The most extensively studied myovirus structure is that of the Gram negative *Escherichia coli* phage, T4^6–11^. The T4 contractile tail shares many features in common with tailocins, although tailocins lack a capsid and do not inject DNA into their target^12, 13^. For CISs, most detailed structural information only covers the initially extended tail conformation. However, contracted state structures have been reported at high resolution for *Pseudomonas* phage E217 and a tailocin, R2 pyocin^14–16^.

Whilst other known myovirus structures share common tail features with T4, E217 and R2 pyocin, the molecular interactions stabilizing their contracted tail state are unknown^16–19^. Moreover, it is not known to what extent the subunit movements within the tail during contraction are the same across myoviruses or CISs in general. Whilst a number of different bacterial cell envelope types are targets for these phages, little is known about the penetration of the thick Gram positive peptidoglycan cell wall; furthermore there has been no explanation of the way phages are able to penetrate the additional protective proteinaceous S-layer found in the majority of eubacteria^20^. An important question is whether S-layer penetrating contractile phages have adaptations to perform the extra work likely to be required to overcome this barrier.

Many bacterial species, including the notable Gram positive pathogen *Clostridioides difficile,* produce a proteinaceous S-layer that forms the outermost surface of the cell envelope. *C. difficile* infection (CDI) is the most common cause of antibiotic-associated diarrhoea, resulting in significant morbidity and mortality in susceptible populations^21^. Phages have been proposed as alternative antimicrobials to treat CDI^22^; naturally occurring phages can be used as a template for the design of such precision antimicrobials^23, 24^. To advance designs, we need to describe the essential structural components and the mechanics involved in binding to the host cell and penetration of the host cell envelope. Here we describe the structure of φCD508, a contractile phage that infects clinically relevant strains of *C. difficile*^25^; its receptor is the major S-layer protein SlpA^24, 26, 27^.

We reveal the high resolution structure of the fully intact virion in both its extended and contracted form. We explain why the extent of contraction is significantly less than that reported for other phage structures. This has important implications for the mechanism of S-layer penetrating phages in general. We also highlight additional and significant key differences to previously characterised myoviruses: the baseplate of the tail is less complex; the needle tip is much more compact than that of other CISs and remarkably, appears to lack enzymatic functions, suggesting that the mode of envelope penetration may be purely mechanical.

## Overall structure of φCD508

φCD508 is a myovirus^25^, with an overall tail length of 225 nm. Structures of *C. difficile* phage φCD508 were determined by single particle EM of frozen hydrated samples (Fig. 1a, c); the resolution ranged from 2.6 Å to 4.0 Å for the extended phage, and 2.9 Å to 4.2 Å for the contracted form (Extended Data Table 1, Extended Data Fig. 1). In the extended state, 14 of the 17 predicted structural proteins identified by mass spectrometry were revealed and modelled at or close to their full chain length. A short section of the tape measure protein (gp59) was also modelled. Each protein chain was built *ab initio* where resolution was sufficient, or was first modelled by RosettaFold^28^ or AlphaFold^29^ and then fitted into the maps of both the extended and contracted phage (Fig. 1b). For additional structural details, refer to the Supplementary Information.

**Figure 1.**
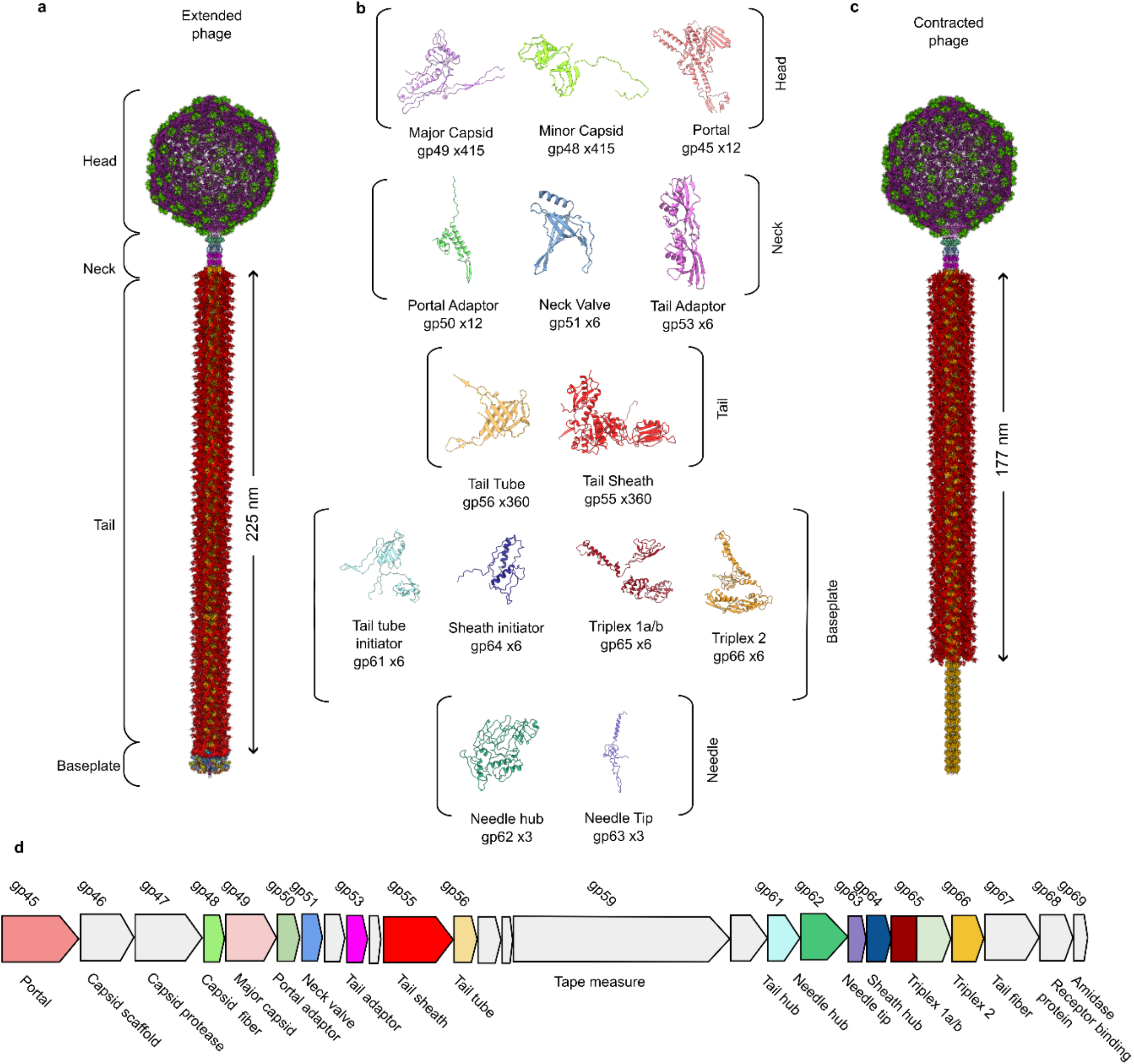
| Structural overview of φCD508. a. Composite model of entire extended φCD508 phage generated from overlapping features of each model. **b.** Gallery of proteins built into cryoEM maps of φCD508, with protein name, gene product number, and copy number in entire extended phage. **b.** Composite model of entire contracted φCD508 phage generated from overlapping features of each model. **d.** Genome organisation of φCD508 structural cassette consisting of gene products 45 to 68. Proteins built into CryoEM maps are coloured as in the gallery, and proteins, where no model could be built, are coloured grey.

The virion can be divided into five distinct regions: the head, neck, tail, baseplate, and needle (Fig. 1a, c). The structural proteins are encoded within a structural cassette (gp45-gp68) of the 74 gene φCD508 genome (Fig. 1d).

The **head** (Fig. 2a) is formed of a near-complete T=7 icosahedral capsid^30^, 650 Å in diameter, and houses the genomic DNA. It is made up of 415 and 420 copies of two capsomer proteins, the major capsid protein gp49 and the capsid decoration protein gp48 respectively (Extended Fig. 2a, b). Within the capsid of the extended phage, internal density shows layers of packaged DNA (Fig. 2a). The contracted virion capsid reconstruction contains an identical arrangement of major capsid protein and capsid decoration protein but lacks internal density indicating that the DNA has been ejected from the head.

**Figure 2.**
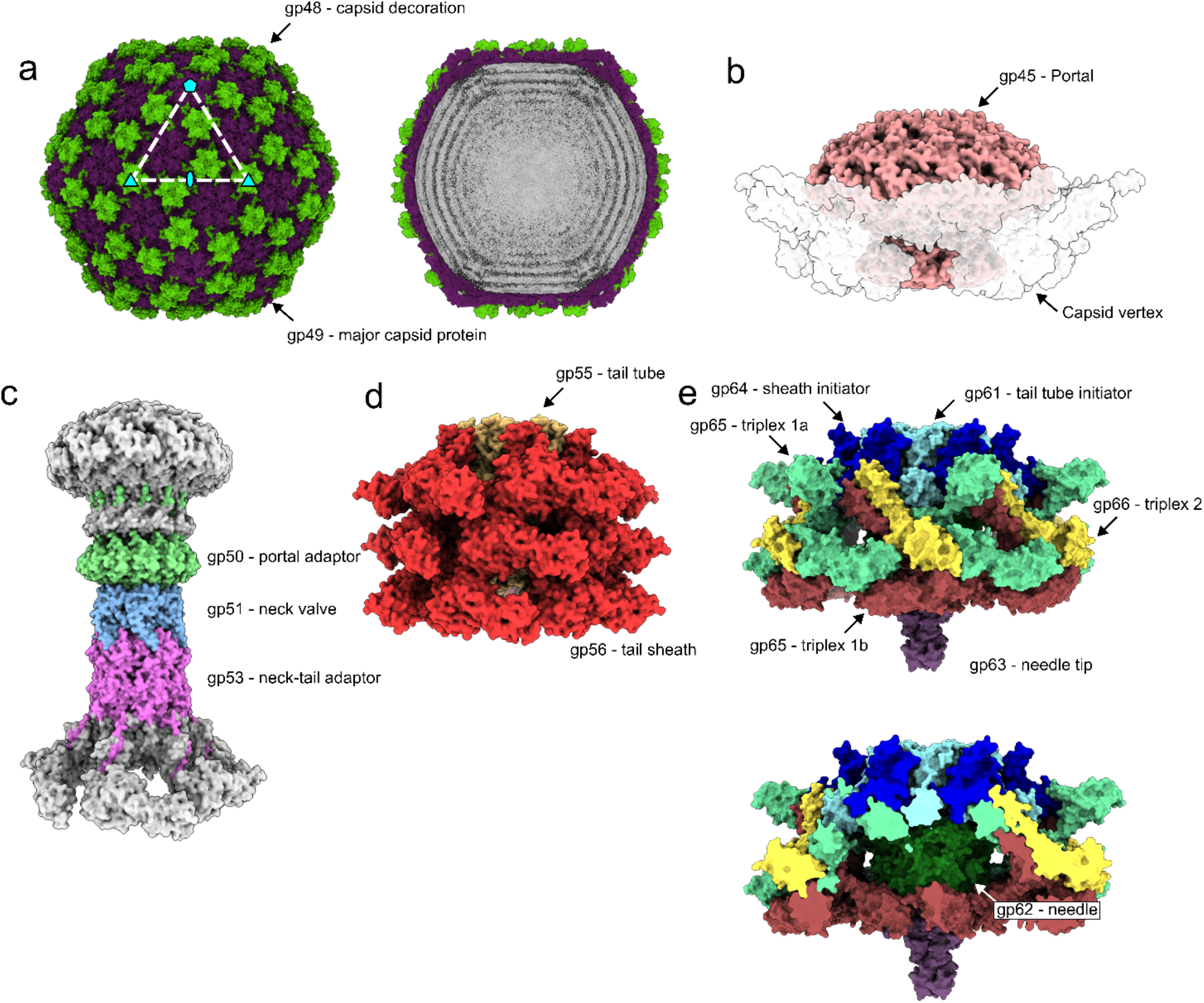
| Structural assemblies of φCD508. Shaded surface representation of φCD508 assemblies as determined by single particle analysis. **a.** Capsid consisting of gp48 and gp49. Slice through extended phage capsid shows DNA layers (right). **b.** Portal protein dodecamer within the unique pentameric capsid vertex, shown with a transparent surface. **c.** Neck proteins shaded by protein identity (portal adaptor = green, neck valve = blue, neck-tail adaptor = pink), with portal and sheath proteins shown in grey for context. **d.** Three layers of sheath (red) and tail tube protein (orange) in the extended state. **e.** Baseplate and needle assembly with hub (blue and cyan), wedge (yellow, mint, and brown), and needle (green and purple).

At the base of the capsid sits a unique opening, centred on what would be a fivefold symmetry axis in a complete icosahedron; it is made up of 12 copies of **portal** protein gp45 (Fig 2b; Extended Data Fig. 2g, h), forming an opening through which DNA is ejected from the capsid. Thus, there is a fivefold to twelvefold symmetry mismatch between the capsid and the portal (Extended Data Fig. 2j, k; Supplementary Information). The portal in turn connects to the **neck** (Fig. 2c; Extended Data Fig. 2i), which is made up of 3 protein complexes with a VIRFAM Type I neck arrangement^31^, comprising one dodecameric ring (gp50) and two hexameric rings (gp51 and gp53) (Fig. 2c). The symmetry is therefore reduced from twelvefold down to sixfold, travelling down the neck. The neck links the capsid to the tail (Fig. 2c).

The first complex in the neck is the **portal adaptor** gp50, which assembles into a twelvefold symmetric ring and acts to resolve a symmetry mismatch between the portal and tail (Fig. 2c, Extended Data Fig. 3a). Gp50 has no structural homology to other known phage neck protein structures (Extended Data Table 2). The **neck valve protein** gp51 forms a hexameric complex and represents the narrowest constriction in the neck lumen, ∼23 Å in diameter at its minimum (Extended Data Fig. 3e). gp51 also has structural homology to other ‘stopper’ proteins such as SPP1 which is proposed to prevent DNA leakage from the capsid^32^. In our structure, strong density in the lumen of the channel weakens around the neck valve (Extended Data Fig. 2i), suggesting the neck valve may also constitute the point at which DNA is prevented from exiting through the tail of the φCD508 prior to contraction. Below the constriction, a weaker density is present which likely constitutes the tape measure protein gp59, although the enforced 6-fold symmetry in the reconstruction prevents identification or building of a model into this region (Extended Data Fig. 2i).

**Figure 3.**
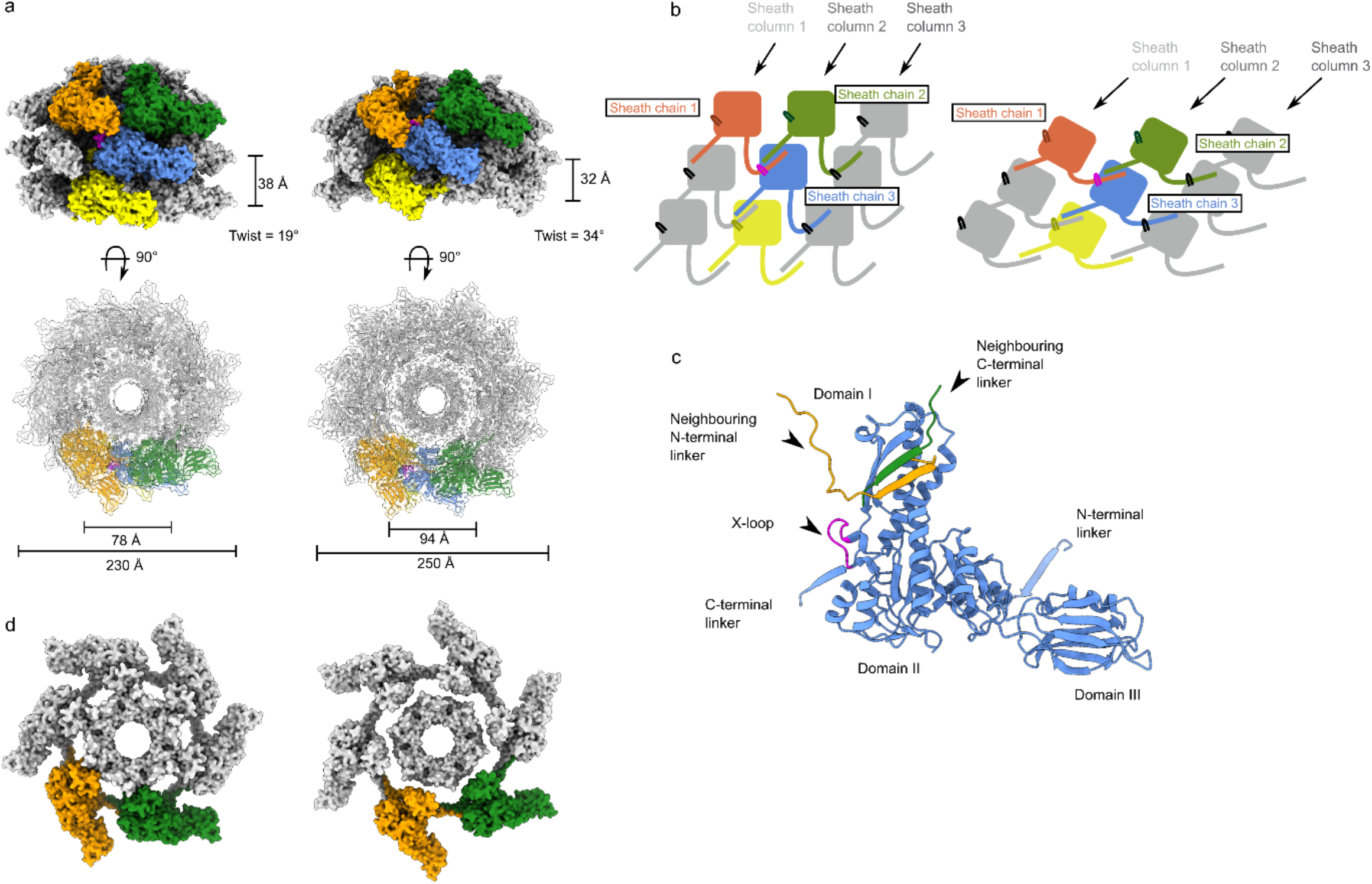
| Reduced contraction of the φCD508 sheath. a. Surface rendering of the extended tail (left) and contracted tail (right), showing helical parameters of sheath proteins and inner and outer sheath dimensions. Four interwoven chains are coloured (orange, green, blue, and yellow), and the X-loop for the blue chain is shown in magenta. **b.** Schematic of mesh network between sheath layers coloured as in a. **c.** Cartoon representation of a single sheath protein showing domain architecture and the X-loop region, coloured as in a. Also shown are the C-terminal linker and N-terminal linker that form the extended mesh between neighbouring chains. **d.** A Surface rendering of the top layer of the sheath and tail tube proteins as in a. In the extended state (left), the sheath forms extensive interactions between the sheath and tail tube. In the contracted form (right), the modelled tail tube is able to pass through the sheath ring.

The **tail** is attached to the neck and made up of a stack of 57 nested hexameric tail protein rings, with an inner ring of **tail tube protein** gp56 and an outer ring of **tail sheath protein** gp55 (Fig. 1, 2d, 3). Each ring is offset from those above and below such that the extended tail assembly can be described as a six-stranded helix, with a twist of 19°, a rise of 39 Å, and an outer diameter of 230 Å (Fig. 1b, 2d, 3a). This arrangement of tail proteins in the extended state is similar to other myovirus and phage tail-like particles. However, a notable feature is that the tail is considerably longer (225 nm) than those of other phages whose structures have been described (Fig. 5).

The main **tail tube** protein, gp56, has the conserved fold seen in other *Caudoviricetes* tail tubes^33–36^. However, gp56 lacks both the ɑ-loop and N-loop seen in other myovirus tail tube proteins, as well as the C-terminal loops seen in siphovirus tail tube proteins^36^ (Extended Data Fig. 4d). This results in a more open packing between rings of the tail tube allowing for bending of the phage in the contracted state (Fig. 4).

**Figure 4.**
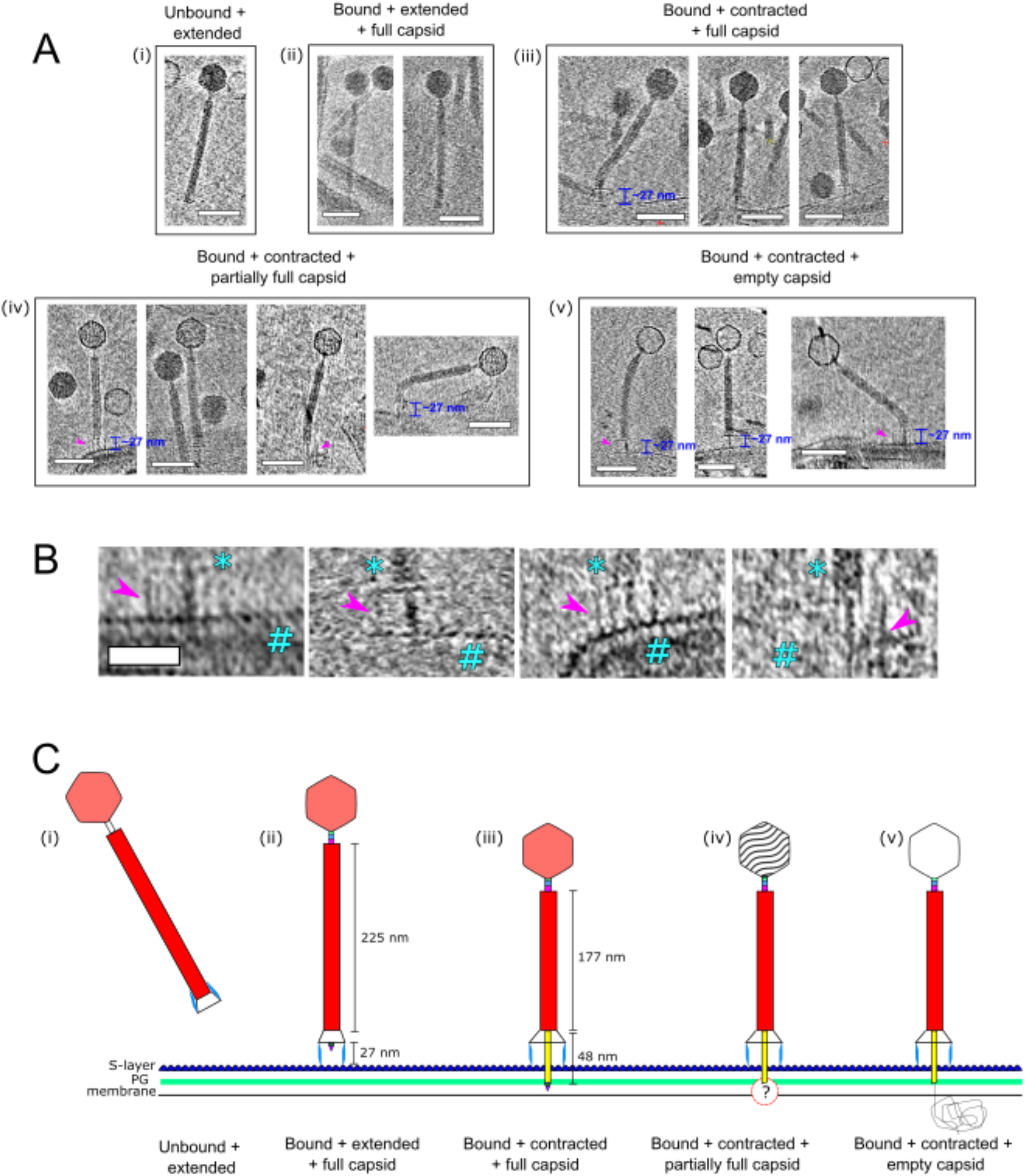
| Model for φCD508 reduced contraction. a. Gallery of φCD508 bacteriophage bound to S-layer fragments from cryoelectron tomograms. The order of images follows the predicted model of phage infection, being free extended phage (i), attachment to the S-layer surface (ii), contraction but no DNA release from the capsid (iii), partial emptying of the capsid (iv), and empty capsids (v). Stages ii-v show examples where the phage is able to bend once contracted. Scale bar = 100 nm **b.** Highlighted baseplates (denoted with *) attached to S-layer fragments (denoted with #), with tail fibers (gp67-gp68) visible and highlighted with magenta arrows. **c.** Schematic model of φCD508 as in a.

The gp55 **tail sheath protein** is made up of three domains (Fig. 3c). The tail tube proximal domain I contains a loop consisting of residues 368 to 378, which is twice as long in φCD508 than in the sheath domains of pyocins, AFP, and PVC (6 residues and 3 residues respectively). For a typical ‘full’ contraction this loop (‘X-loop’) would interact with the linker of the N-terminal β-strand insertion of a neighbouring sheath protein (Fig. 3b, c), significantly reducing the possible range of motion of the linker compared to other known CIS structures (Extended Data Fig. 4e, f).

The tail is terminated by the **baseplate**. Importantly, the baseplate of φCD508 must be adapted for binding and penetration of the host cell S-layer. The baseplate is minimal compared with other structurally characterised phages^11, 17, 18^ consisting of only 6 unique proteins (Fig. 2e). These are organised into a **hub** (Extended Data Fig. 6a) which connects to the tail and a **wedge** which forms a collar around the needle (Fig. 2e, Extended Data Fig. 6c). The baseplate is also expected to include the host attachment proteins gp67 and gp68, constituting the tail fibre and receptor binding proteins respectively, and which are essential for S-layer attachment^27, 37^. Although these could not be sufficiently resolved in the EM maps for atomic modelling, low pass filtered maps show protrusions for the tail fibres (Extended Data Fig. 6f). Within the ring formed by the baseplate proteins lies the **needle** assembly; this is the other essential component presumed to be adapted for S-layer penetration, formed of two proteins, gp62 and gp63 providing a pointed end terminating the tail (Fig. 2e, Extended Data Fig. 6h). It is striking that we find no putative enzymatic domains within the baseplate/needle complex, unlike other comparable assemblies^11, 17, 38^. In urea contracted phage, we observe that the baseplate is lost from the tail, and it could not be resolved even at low resolution. Using the extended phage baseplate, we modelled what the contracted baseplate would look like. One spoke of the baseplate including the baseplate proximal sheath layer, baseplate hub and triplex proteins were aligned onto the contracted sheath. The rigid motion of these proteins would be sufficient to break the dimerisation interactions in the triplex proteins, as well as interactions between the triplex wing domains and the needle protein (Extended Fig. 6g). Breaking these contacts would be sufficient to allow clearance for the tail tube to pass through the baseplate.

## The φCD508 sheath does not contract as much as other CISs

The overall RMSD between all proteins of the neck in the extended state compared to the contracted state is 0.34 Å, indicating that no large conformational change occurs in this region during phage contraction. After contraction and genome release, the neck-proximal tail sheath protein (gp55) remains attached to the neck-tail adaptor gp53 via its C-terminal insertion domain (Extended Data Fig. 3h, i). The neck-proximal sheath protein rotates to accommodate the contraction induced movements of the other sheath proteins in the tail below, pivoting around Gly^260^ in the gp53 C-terminal linker (Extended Data Fig. 3i, k). The terminal sheath ring still maintains some contact with the tail tube. This reduced rotation is limited to the proximal layer, with the next layer below fully dissociating from the tail tube, similar to the arrangement observed for the main body of the contracted tail reconstruction.

Contraction of φCD508 in urea reduces the tail length by just 20% whereas most CIS tails that have been described contract to ∼50% of their extended length; the outer diameter of the sheath increases from 230 Å to 250 Å, and the inner diameter from 78 Å to 94 Å, sufficient to break contacts between gp55 sheath domain I helices and the tail tube gp56 (Fig. 3d). The contracted sheath retains the arrangement of gp55 subunits in a six-stranded helix, but with a decrease in helical rise from 39 Å to 32 Å, and a concomitant increase in twist from 19° to 34°. The contraction induced movement of the gp55 sheath proteins can be modelled as a rigid body pivoting motion about domain I (Extended Data Movie 1, Fig. 3a, b).

Electron cryotomography of φCD508 virions incubated with S-layer fragments confirmed the 20% reduction in sheath length in the presence of the natural receptor (225 nm to 177 nm) (Fig. 4a). In the urea contracted form of φCD508 the baseplate was lost, whereas the baseplate was retained when bound to the native S-layer receptor (Fig. 4b). Notably, at intermediate time points in the incubation we found that some fully contracted virions had not yet released their genome from the capsid (Fig. 4a). This was previously observed in images of the *Staphylococcus aureus* phage phi812^18^.

In both the extended and contracted form, the sheath ring retains integrity through its mesh network, similar to the sheath of other CISs^12–14^ (Fig. 3b, d; Extended Data Movie 1). The overall RMSD between sheath protein monomers in the extended and contracted form is 0.63 Å, confirming a rigid body movement of these proteins. In helical reconstructions of contracted tail, the tail tube cannot be resolved due to a mismatch between the helical parameters of the sheath and tail tube. As the tail tube does not contract, the resulting mismatch in length drives the tail tube through the bottom of the baseplate. The measured 20% reduction (48 nm) in tail length is consistent with a decrease in helical rise from 39 Å to 32 Å for 57 rings of sheath protein.

In the contracted phage tail, which lacks the packing interactions between the tail tube and the sheath, the bending stiffness (persistence length) is significantly reduced (Fig. 4a). Compared to other known contracted CIS structures, the more open packing of subunits both within the tail tube and tail sheath (Fig. 3, Extended Data Fig. 4) may allow the tail to be more flexible similar to the intermediate states of A511 contraction where flexibility is also observed^17^.

## Discussion

Here we present a complete atomic model of the contractile bacteriophage, φCD508, in both extended and contracted states (Fig. 1). φCD508 is significant as it shows adaptations for binding to and penetration of the host cell S-layer, whereas all of the phage that have been structurally characterised previously, infect bacterial hosts that lack this widely encountered barrier^6, 15, 17–19, 35^. The most striking difference between φCD508 and other known CIS structures is the much smaller relative contraction (Fig. 1, 5). There are other phages described in the literature for which reduced contraction has been observed, but not yet analysed^39–41^. Remarkably, these phages also infect Gram positive species that have S-layers. It is also notable that the tail tube protrusion length in these phages seems independent of total tail length (Fig. 5). This could represent an adaptation towards hosts with S-layers in general. To the best of our knowledge, for all other CISs studied in detail the tail tube protrusion length is correlated with the total tail length (Fig. 5). The other remarkable feature of φCD508 is its minimal baseplate with compact needle tip, and no enzymatic domains.

**Figure 5.**
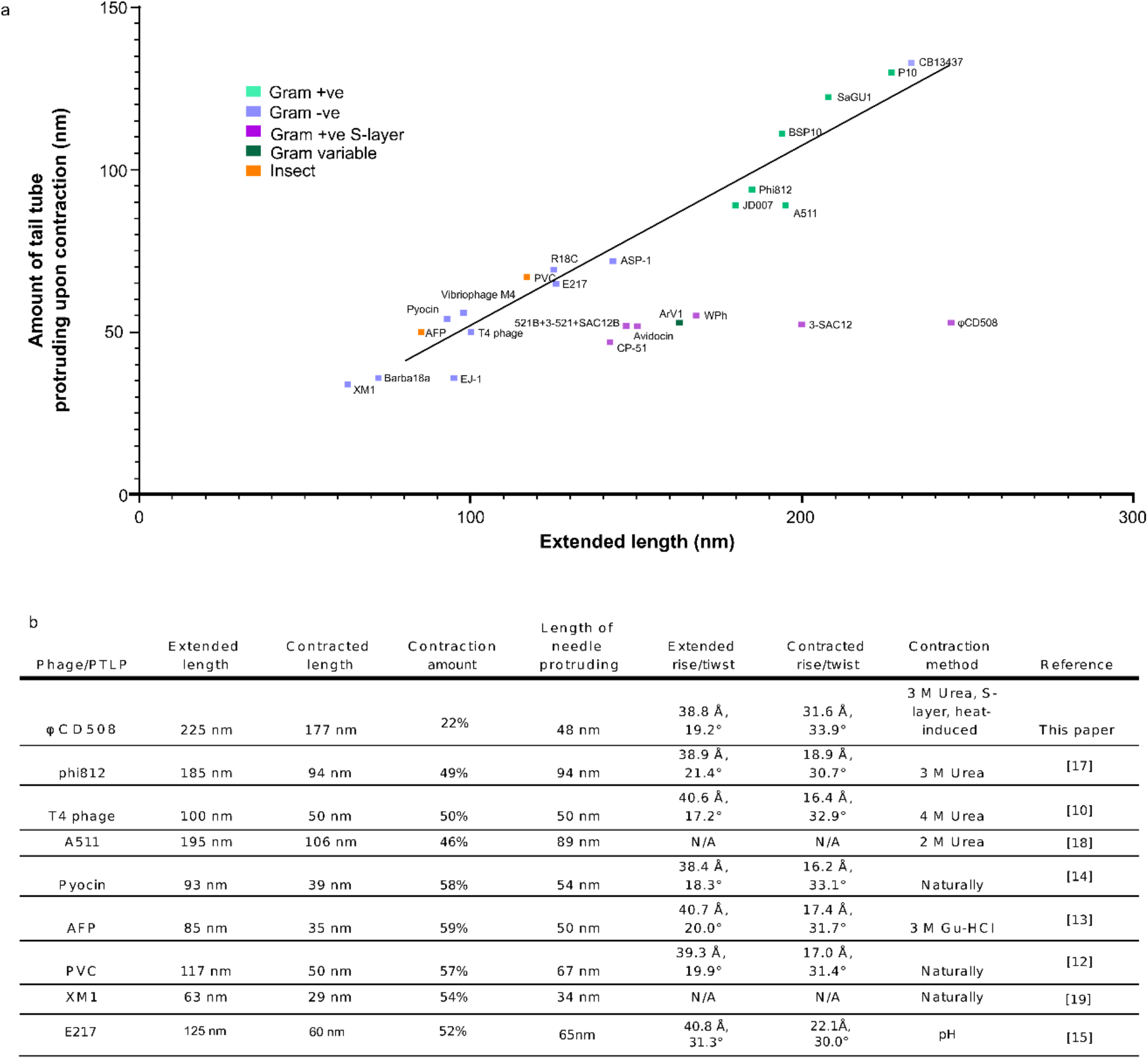
| Comparson of contraction parameters in CISs. a. Graph of the extended tail length vs. the amount of tail tube protruding upon contraction for a number of contractile bacteriophage and phage tail-like particles (PTLPs) from literature. Each particle is coloured based on the bacterial/insect host type. ‘Conventional’ virions fit well along a trendline (R^2^ = 0.975). **b.** A table of tail parameters for bacteriophage and PTLPs for which structural data are available.

We reason that the reduced contraction of φCD508 is mediated by the sheath, as phages in which the baseplate has become detached remain in the same contracted state (Fig. 4), suggesting this state is stabilised by the sheath proteins alone and not influenced by partial rearrangement in the baseplate. The N-terminal linker forms part of the mesh that connects the sheath proteins to one another, and has been shown to act as a hinge maintaining links between subunits through the sheath contraction^42, 43^. The increased length of the X-loop acts to restrict the motion of this N-terminal hinge compared with other CISs (Fig. 3c; Extended Data Fig. 4e). Further, the X-loop blocks complete contraction of the tail for φCD508 (Fig. 3); we modelled the φCD508 sheath proteins into a more contracted conformation, based on pyocin trunk structures^43^, but increased contraction of φCD508 sheath proteins led to steric clashes between the X-loop and the neighbouring N-terminal linker as well as domain II of a neighbouring sheath (Extended Data Fig. 4f).

The reduced compression between layers of sheath protein is not associated with a reduction in rotation. The helical twist between sheath disc layers increases from 19° to 34° during contraction, similar to other CISs^12, 13, 17, 18, 43^ (Fig. 5). A recent study showed that the rotation and translation of the tail sheath of pyocins occurs simultaneously during contraction^42^. Therefore, further contraction of φCD508 would lead to an over-rotation that cannot be accommodated by the mesh-like interactions between sheath proteins.

The disconnect between tail tube protrusion length and overall tail length in phages that infect S-layer coated bacteria is striking (Fig. 5) and may reflect the different energetics involved in penetrating the very diverse types of S-layers found in different species^20^, the need to accommodate different functionalities in the tape measure protein and the degree to which enzymatic attack contributes to the penetration process. For example, the phage tail-like Avidocins that also target *C. difficile* strains, have shorter tails than φCD508^44^ but also incorporate predicted hydrolytic enzymes (Fig. 5).

Uniquely, here we were able to visualize exposure of the tail tube in the context of S-layer penetration. The major S-layer protein, SlpA, acts as the φCD508 receptor; we have previously shown that sequence variation within the main S-layer subunit SlpA (S-layer cassette type, SLCT) correlates with phage infection spectrum and that heterologous expression of the SLCT-10 SlpA alone was sufficient to sensitise a normally resistant *C. difficile* strain to φCD508^27^. Tomograms of φCD508 bound to S-layer fragments (Fig. 4a, b) show how the phage interacts with the host during infection. Side views of φCD508 bound to the fragments show that the baseplate is held ∼27 nm from the S-layer surface by the tail fibres (Fig. 4a, b). The thickness of the *C. difficile* cellular envelope is ∼37 nm, and so combined with the ∼27 nm distance from the S-layer, 48 nm of exposed tail tube would appear to be insufficient to reach the cell membrane and enter the cytoplasm. In T4 phage infection, the membrane is actively pinched outwards by 16 nm normal to its plane to meet the tail tube. It has been proposed that the tape measure protein gp29, containing a hydrophobic stretch, plays a role in this pinching to form a channel through which DNA is injected^45^. As our S-layer fragments lack membrane (Fig. 4), we cannot test for a similar pinching phenomenon, however, the φCD508 tape measure protein gp59 contains at least three predicted transmembrane stretches and may play a similar role to T4 gp29 (Extended Data Fig. 8). We also imaged phages contracted on whole cells; although the cells were too thick to visualise the membrane state upon phage adsorption and contraction, we did not observe any additional contraction compared with phages contracted on S-layer fragments. This suggests indeed that the membrane must be drawn closer to the end of the tail tube by some as yet unidentified mechanism.

A search of genome sequences of phages infecting bacteria with S-layers suggests that the particularly compact φCD508-form of the needle, may represent a widespread, but previously unreported adaptation (Extended Data Fig. 9b, c). The terminus of the φCD508 needle is indeed an apex domain as already reported in other phages^46^; it displays a conserved hexahistidine motif coordinating a putative iron ion (Extended Fig. 6, 7). However, the notable feature of φCD508, and the other S-layer penetrating phages we have analysed, is that the needle lacks the otherwise widely conserved β-helix (Extended Data Fig. 9a). This β-helix has been implicated in piercing the outer membrane of Gram negative bacteria^46^, but is also found widely in CISs targeting Gram positive bacteria and eukaryotes^13, 46, 47^. The compact structure of the φCD508 type of needle is perhaps an adaptation towards easing it across the S-layer barrier, although the role of the needle apex itself in infection remains unclear^48^.

We propose a model for φCD508 phage infection (Fig. 4c). Initially, the phage binds to SlpA on the surface of the *C. difficile* cell^49^ via receptor binding domains on the tail fibres. It is likely that binding is initially reversible and that the phage conducts a ‘random walk’ across the host cell surface (Fig. 4c (i), (ii)). These initially transient interactions have been observed or proposed for a number of other phages including those infecting *C. difficile*^50–53^. Only when the phage is committed to irreversible binding to the SlpA receptor does a structural change take place in the baseplate leading to tail sheath contraction, but only by 20% (Fig. 4c (ii), (iii)). It is tempting to speculate that the contraction drives the compact needle proteins through the surface of the cell envelope. However, the details of how this would happen are far from clear. Firstly, we do not know if the needle is driven through the very closely packed SlpA lattice itself or through one of the lattice defects/grain boundaries in the natural S-layer^26^; the latter might require less force but how weak points in the S-layer could be detected is unknown.

Once the S-layer barrier has been breached, the cell wall must then be penetrated. It is remarkable that φCD508 appears to have no enzymatic means of doing this, so that it would seem that mechanical work must be performed at this step. The energetics of this remain to be determined although it should be noted that the cell wall is permeated by pores and cavities that may be weak points^54^.

The final barrier is the cell membrane. As the contraction length is insufficient for the tail tube to reach the cytoplasm, the membrane may bulge outwards to meet the tail tube in a process analogous to T4 phage infection^45^. The compressed DNA in the capsid might expel the trimeric tape measure protein gp59, which then forms a membrane channel via the predicted transmembrane helices of each polypeptide and through which DNA can be injected or drawn (Extended Data Fig. 8). It is also possible that local damage to the cell wall results in the membrane bulging as a consequence of the cell turgor pressure.

The detailed mechanism by which the potential energy of the extended state is converted into work to penetrate the host envelope is unknown for most CISs. Imaging of contraction intermediates of phages indicates that contraction is triggered at the baseplate and propagates as a wave along the tail^17, 55, 56^. It has been proposed that long-lived intermediates represent a stage in infection of Gram positive bacteria at which digestion of the cell wall is carried out by enzymes located at the tail tube tip^17^. In φCD508 we have not identified any intermediates in which the tail region proximal to the head remains in the uncontracted conformation^17^. This is consistent with the hypothesis that penetration of the S-layer and cell wall does not involve any enzymatic activity.

Fraser *et al.* modelled and experimentally measured the energetics and forces generated through the contraction of an R-type pyocin^42^. Interfacial sheath-sheath subunit interactions dominated the energetics; the free energy difference between the extended and contracted states was found to be largely enthalpic. The model suggests that because of the wavelike nature of the contraction, CISs that are longer than the observed ‘contraction wavelength’ generate similar forces. This raises the question of why the tail of φCD508 is relatively long and why, although the sheath-sheath rearrangement is not as extensive as that in other CISs, sufficient force is still generated. It is possible that the ‘contraction wavelength’^42^ is longer in φCD508 so that force is maintained over a longer distance. Moreover, the φCD508 tail becomes very flexible in the contracted state (Fig. 4a) possibly adding an entropic component to the total free energy of contraction, thus compensating for the relatively smaller sheath-sheath interfacial rearrangement.

It is clear that φCD508 is a member of a novel class of contractile tail phages with reduced contraction (Fig. 5), including phages infecting other S-layer producing bacteria. Our CryoEM structures show that although many common features are shared between φCD508 and other CIS structures, there are some very significant differences. In order to develop phages as novel antimicrobials, and to engineer phages targeting clinically important strains of *C. difficile*, characterisation of the advantage these adaptations confer and the detailed interactions with SlpA will be vital.

## Data Availability

CryoEM maps will be deposited with the Electron Microscopy Data Bank and coordinates will be deposited with the Protein Data Bank. The complete φCD508 genome sequence is available through the National Center for Biotechnology Information database under the accession number OR295560.

## Acknowledgements

We thank Svetomir Tzokov of the Faculty of Science Electron Microscopy Facility, University of Sheffield, for assistance with EM and Adelina Acosta Martin of the Faculty of Science Mass Spectrometry Facility for assistance with MS analysis. We acknowledge Diamond Light Source for access and support of the cryo-EM facilities at the UK national electron Bio-Imaging Centre (eBIC), proposal EM19832, funded by the Wellcome Trust, MRC and BBSRC, Yun Song and Vinod Kumar Vogirala for help with data collection and Mathew Arnold for help with data processing. We are most grateful for helpful discussions with Rebekkah Menday, David Rice, Rebecca Corrigan and Indrajit Lahiri.

This work was funded by BBSRC grant BB/P02002X/1 to PAB and RPF and The University of Sheffield’s Imagine:Imaging Life programme.

For the purpose of open access, the authors have applied a Creative Commons Attribution (CC BY) licence to any Author Accepted Manuscript version arising.

## Author contributions

JSW designed the study, carried out all experiments, collected and analysed data, wrote and revised the manuscript. LCF designed experiments, analysed data, and revised the manuscript. RPF designed the study, analysed data, supervised the study, wrote and revised the manuscript. PAB designed the study, analysed and interpreted EM data, supervised the study, wrote and revised the manuscript.

## Materials and correspondence

Requests should be addressed to Per Bullough or Robert Fagan.

