## Supplemental text and figures for "Molecular mechanism of bacteriophage tail contraction-structure of an S-layer-penetrating bacteriophage"

**Supplementary Information**

### Methods

#### Phage production and purification

φCD508 phage were propagated in *C. difficile* strain CD117. Cells from overnight culture were inoculated into 200 ml TY media and grown to an $OD_{600nm}$ of 0.1, before addition of 10 mM $MgCl_2$, 10 mM $CaCl_2$, and φCD508 stock to a multiplicity of infection (M.O.I) of <0.1. Cells were grown for ~4 hr, by which time most of the cells were lysed by the phage. The cells were then harvested at 4000 x g and the supernatant filtered through 0.45 µm filters.

The filtered supernatant was centrifuged at 69,000 x g for 2 hours at 4 °C to pellet the phage, and then resuspended in 2 ml 50 mM HEPES pH 7.4, 10 mM $CaCl_2$, 10 mM $MgCl_2$, and 10 mM NaCl, overnight with rotation at 4 °C. The resuspended phage were mixed with 1 ml chloroform and centrifuged at 3,000 x g for 10 min. The soluble fractions were overlaid on a CsCl gradient with 1 ml each of 1.45 g/ml, 1.5 g/ml, and 1.7 g/ml CsCl. These were centrifuged at 152,000 x g for 4 hr at 15 °C. A white band corresponding to φCD508 was extracted by syringe and dialysed against 500 ml 50 mM HEPES pH 7.5, 10 mM NaCl, 10 mM $MgCl_2$ and 10 mM $CaCl_2$ overnight at 4 °C. Samples were stored at 4 °C.

#### Phage urea contraction

500 µl φCD508 at 6 mg/ml was diluted to 10 ml in filtered 50 mM HEPES pH 7.4, 10 mM $CaCl_2$, 10 mM $MgCl_2$, 10 mM NaCl, 3 M urea and incubated at 4 °C for 2 hr. 30 µg/ml DNaseI was added and incubated at 17 °C for 10 min, made up to 30 ml in 50 mM HEPES pH 7.4, 10 mM $CaCl_2$, 10 mM $MgCl_2$, 10 mM NaCl, and centrifuged at 74,500 x g for 1 hr at 4 °C. The pellet was resuspended in 70 µl milliQ water; EM grids were prepared immediately.

**CryoEM data collection and image processing**

Extended phage at 2 mg/ml were vitrified by double blotting onto Quantifoil R2/2 grids that had been glow discharged (Cressington 208C, Cressington, Watford, England) for 10 s. Contracted phage at 1.4 mg/ml were vitrified by single blotting onto Quantifoil R2/2 grids that had been glow discharged for 14 s. For both datasets, micrographs were collected on a FEI Krios Titan operated at 300 kV and fitted with a Gatan K3 camera. A nominal magnification of 81,000x was used in super-resolution mode, with a super-resolution pixel size of 0.58 Å/pixel. 14,533 movies were collected for extended phage, and 9,557 movies were collected for contracted phage, each containing 40 frames and with a total dose of 42 e/Å$^2$. Raw movies were motion corrected using MotionCor2[1] with 2x binning, to give a nominal pixel size of 1.06 Å/pixel, and defocus values were estimated using ctffind 4.1[2].

**Helical reconstruction of tail tube and sheath in extended conformation**

Extended phage tails were picked automatically using crYOLO[3]. 344 helical segments were picked from 64 extended phage micrographs using EMAN helix boxer[4], and then used to train the crYOLO picking model. The model was used to pick 123,058 helical segments from 14,533 micrographs with a picking threshold of 0.15. Helix start and end coordinates were used to extract 500 pixel boxes in RELION[5], with a helical rise of 40 Å. Helical rise and twist were estimated by first performing 3D reconstruction with no helical symmetry. The resulting map went to 7.3 Å resolution, with clear secondary structure features present. This was used to ascertain helical parameters, and helical reconstruction was then performed in cryoSPARC[6], converging at 2.7 Å resolution with a helical twist of 19.2° and rise of 38.9 Å.

Contracted phage tails were picked using cryoSPARC filament tracer, with a filament diameter of 210 Å and separation distance between segments of 0.5 diameters to prevent overlapping segments. 151,339 particles were extracted with a box size of 580 pixels and 2D classification was used to select good segments, with 100 classes. 17 good classes containing 128,963 particles were re-extracted with a 400 pixel box size, and used for *ab initio* reconstruction in C1. Helical parameters were estimated from a homogeneous refinement as 33.9° twist and 31.6 Å rise, which were subsequently used in helix refinement enforcing C6 symmetry and using non-uniform refinement. Refinement converged at 4.2 Å resolution with a 34.0° twist and 31.8 Å rise

**Capsid reconstruction**

Extended phage capsids were picked automatically using crYOLO, after training on 30 micrographs. 2D classification was used to select for classes with high resolution secondary structure features, and capsids that contained DNA. ~70K particles were selected, and the coordinates used to extract 1000 pixel boxes in RELION. An initial model was generated by *ab initio* reconstruction in cryoSPARC, enforcing I1 symmetry. The initial model was then used in 3D refinement with I1 symmetry.

Contracted phage capsids were picked using RELION autopicker with 0.4 picking threshold. 42,410 particle coordinates were used to extract particles with 1000 pixel box size. *Ab initio* reconstruction was performed with 10,000 particles and enforced C5 symmetry, and used as a reference in homogeneous refinement with I symmetry. Refinement converged at 3.3 Å resolution.

**Baseplate reconstruction**

Baseplates from the extended phage dataset were picked using crYOLO after training on a subset of 139 particles from 56 micrographs; 23,587 particles were extracted with 500 pixel box sizes, and were subjected to 25 iterations of 2D classification in RELION with 30 classes and 480 Å mask. Good classes containing secondary structural features for the baseplate were selected, and 4,281 particles from these classes were used for an initial reconstruction. 50 equally-spaced templates were generated from the initial reconstruction in CryoSPARC and used to pick the final particle set. 2D classes containing baseplates were selected, incorporating 19,276 particles. A C6 *ab initio* reconstruction was used as reference for homogeneous refinement imposing C6 symmetry, followed by local CTF refinement and non-uniform refinement. The final reconstruction converged at 3.4 Å, and was further sharpened with a B-factor of -64.7 for inspection and model building.

**Portal and neck reconstruction**

In order to locate the extended portal and neck vertex, an I4 icosahedral capsid reconstruction was generated in RELION to align a 5-fold vertex along the z axis. These I4 aligned particles were symmetry expanded using *relion_particle_symmetry_expand --i run_data.star --sym I4 --o expanded_particles.star*. Expanded particles were then re-extracted with a 1000 pixel box size rescaled to 126 pixels, and then used in focused 3D classification with a tight cylindrical mask and a T-number (tau fudge) of 20. Two classes containing 230,382 particles had density for the neck and portal, and were re-extracted with recentring offset of 37 pixels in the z direction, to centre on the portal vertex. Duplicate particles were removed from the recentred particles, with a minimum distance cut-off of 108 Å. 70,554 remaining particles were re-extracted with a box size of 300 pixels and run through 2D classification in CryoSPARC. 38,968 particles containing clear secondary structure features were used in *ab initio* reconstruction with C12 symmetry, followed by refinement in CryoSPARC, again with C12 symmetry, and with 3 final passes after convergence; this improved the quality of the density, converging at 3.3 Å. Local CTF refinement was run in cryoSPARC, and homogeneous refinement run with the CTF refined particles, converging at 2.6 Å.

The extended phage neck was reconstructed in a similar way to the portal, by re-extracting the classes containing portal density from focused 3D classification, but with an offset of 50 pixels in the z-direction, which centred the neck proteins in the extracted boxes and applying C6 symmetry in both *ab initio* reconstruction and homogeneous refinement of the neck in cryoSPARC. Refinement converged at 3.8 Å, and then 3.4 Å after local CTF refinement.

Contracted phage portals and necks were generated in the same way. An I4 capsid reconstruction from 33,824 autopicked capsids was symmetry expanded and 1000 pixel boxes were rescaled to 126 pixels. 3D classification was run with 10 classes, a tight cylindrical mask used for the extended portal localisation, and a T-number of 20. One class containing 116,487 particles was extracted with an offset of 37 pixels, and duplicates were removed with a minimum overlap distance of 60 Å, leaving 31,079 particles. These particles were extracted with a box size of 300 pixels and were imported to cryoSPARC. 2D classification was run with 50 classes, and 12 classes containing clear portal density were selected, containing 26,404 particles. The C12 reconstruction was generated using an *ab initio* model with enforced C12 symmetry, and converged at 2.9 Å; the C5 portal reconstruction was generated using the C12 *ab initio* model but enforcing C5 symmetry during homogenous refinement, converging at 3.5 Å.

Overlapping features in the C12 portal and C6 neck reconstructions were used to generate the composite atomic model after building.

**Portal / Capsid mismatch reconstruction**

Firstly, aligned particles from the C12 portal reconstruction were used in homogenous refinement in cryoSPARC using C5 symmetry, with a C5 enforced *ab initio* model as a reference. This reconstruction resolved the capsid density surrounding the portal complex. Next, the resulting particles and orientations were exported to RELION using the UCSF pyem routine csparc2star.py[7], and symmetry expanded with C5 symmetry. The expanded particle sets were used in 3D classification with a mask around the portal, without alignment and with no symmetry imposed. Classification resulted in five classes containing ~20% of particles in each class. One class was chosen and further refined by 3D refinement. A mask was generated using Chimera's segment function to mask the capsid and portal complex, for masked refinement and post processing[8].

**Model building, refinement, and validation.**

Purified phage were analysed by LC-MS/MS and identified structural proteins were built *ab initio* where resolution was sufficient, or first modelled by RosettaFold[9] or AlphaFold[10], and then fitted into the maps of both the extended and contracted phage. Model building was done *ab initio* using Coot[11] where the resolution permitted. A poly-Ala trace was generated and then side chains were added in Phenix[12] using sequence from map, with candidates from mass spectrometry data and genome data to guide identification. Where matches were found, the sequence was completed and further refined in Phenix with real space refinement, whilst manually correcting errors with Isolde's model rebuilding tools[13]. Validation was performed using MolProbity[14].

Density for the distal domain of the sheath protein gp55 was too poor to build *ab initio*, and so the gp55 structure was also predicted by Robetta[15], and compared with the initially built proximal domains; a fit of 0.9 Å RMSD was achieved. The predicted distal domain was fitted with restrained flexible fitting into the map using Isolde.

Baseplate proteins gp65 and gp66 were also built partially from AlphaFold predicted structures using the colabfold build of AlphaFold[16], by first building *ab initio* through interpretable density, and then fitting of predicted structures and flexible fitting of the remaining domains. Completed models were run through Isolde with restrained flexible fitting to fit main chain and side chains into maps of different sharpening B-factors.

**φCD508 ghost tomography**

*C. difficile* strain CD117 S-layer ghosts were generated as described previously[17]. Briefly, 80 ml TY was inoculated with CD117 to 0.05 OD and grown to OD 0.7. Cultures were centrifuged at 4 °C for 15 mins at 2000 x g, then resuspended in 30 ml ice cold deionised water. This was repeated with a resuspension in 15 ml. This was added to 15 ml pre-cooled 212 – 300 µm acid-washed glass beads (Sigma) on ice in a homogenisation flask and homogenised for 30s, then placed on ice for 5 mins. The supernatant was decanted and centrifuged at 800 x g for 10 mins at 4 °C and the supernatant again centrifuged at 3000 x g for 10 mins at 4 °C. The pellet was resuspended in 400 µl ice cold 1M NaCl and centrifuged for 10 mins at 4 °C. Finally, the pellet was resuspended in 200 µl phage buffer (50 mM NaCl) and flash frozen in liquid nitrogen in PCR tubes in 25 µl aliquots. 2.7 mg/ml CD508 and S-layer suspension were mixed 1:1 (vol:vol), and incubated at 37 °C for 5 or 30 min and then placed on ice. 8 µl BSA-treated gold fiducials[18] were mixed with 20 µl phage/S-layer mix, and 3.5 µl sample was applied to R3.5/1 Quantifoil grids that had been glow discharged for 14 s. Sample was adsorbed for 1 min and then blotted for 4 s using a Leica GP2 blotting device before plunging into liquid ethane.

Tomograms were collected with Tomo software on a Tecnai Arctica 200 kV microscope fitted with a Falcon III detector. Tilt series were collected from -60° to +60° with a 3° tilt increment, starting from 20° and sweeping through to -60° before completing from 20° to 60°. At each tilt angle, images were collected with 1.2 e/Å$^2$ over 10 frames. Tomograms were collected with an applied defocus ranging from -3 to -8 µm. Tomograms were reconstructed using IMOD using gold beads as fiducials for generating an alignment model[19]. Tomograms were reconstructed with weight back projection, and with 15 iterations of SIRT-like filtering and 2-3 x binning in order to measure phage lengths.

**φCD508 tomogram tail measurements**

Reconstructed tomograms were used to measure tail lengths in IMOD by selecting the neck proximal and baseplate proximal sheath layers, and using analyse tubes in IMOD to calculate lengths. Virions with full capsids, partially emptied capsids, and emptied capsids were measured and used to determine average lengths.

**Phage genome sequencing**

Genomic DNA from φCD508 was extracted by phenol-chloroform as described previously[20], then further purified using Agencourt AMPure XP magnetic beads (Beckman Coulter). Phage DNA libraries were prepared for Illumina sequencing using the NEBNext® Ultra™ II FS DNA Library Prep Kit for Illumina® (New England BioLabs) according to the manufacturer’s instructions. Sequencing was performed on an Illumina Nextseq500 sequencer (paired end 2 x 75 bp) at the RNomic Platform of the Université de Sherbrooke (Sherbrooke, Québec, Canada). Trimming of the raw reads was performed using fastp (v.0.20.0)[21]. Clean reads were then assembled *de novo* using SpaDes (v.3.14.0)[22] with default options. A single contig of 49,272 bp with a coverage of 1,232x was obtained.

**Genome annotation**

Genome annotation was performed using PROKKA (v.1.14.6)[23] run locally with the vCONTACT2 protein sequence database downloaded on March 31, 2023, using the INPHARED perl script[24]. The e-value threshold was set at 10-3. Structural proteins for which a function could not be assigned using PROKKA but that were detected by mass spectrometry were annotated based on the function predicted from the structure reconstruction. The genomic map was created using Benchling v.2.1.2 and finalised with Inkscape v.1.2.1.

### Supplementary Discussion

#### *Structure of the capsid and portal*

The **capsid** from extended phages (Fig. 2a) was reconstructed with icosahedral symmetry imposed, to 3.4 Å resolution; it consists of two proteins, the major capsid protein gp49 (Extended Data Fig. 2a), and the capsid decoration protein gp48 (Extended Data Fig. 2b). The major capsid protein assembles to form hexameric faces and pentameric vertices in a T=7 *laevo* icosahedral arrangement (Extended Data Fig. 2d)[25]. Thus, the icosahedral asymmetric unit consists of 7 gp49 polypeptides and 7 gp48 polypeptides (Extended Data Fig. 2c); 6 gp49 subunits are arranged around a quasi-sixfold symmetric axis with the seventh arranged with four additional partners around an icosahedral fivefold symmetry axis. The gp48 polypeptides associate into trimers; two trimers assembled around a quasi-symmetric threefold axis and one polypeptide assembled with partners around a true icosahedral threefold symmetry axis (Extended Data Fig. 2a). The major capsid protein gp49 adopts the typical HK97 fold[26], consisting of an axial domain, a peripheral domain, an extended loop, and an N-terminal arm (Extended Data Fig. 2a). The extended loops point towards the centre of the hexameric face (Extended Data Fig. 2c). In addition, a small insertion domain formed from residues 209-231 (Extended Data Fig. 2a) forms a β-hairpin that associates with βB of a neighbouring capsid decoration protein, contributing to capsid stability.

The capsid decoration protein gp48 forms trimeric protrusions from the capsid surface (Fig. 2, Extended Data Fig. 2c, d). The decoration protein has structural homology to siphovirus TW1 and podovirus φ29 capsid decoration proteins[27,28] (Extended Data Table 2), but lacks the long extension present at the C-terminus of the φ29 protein. Each gp48 trimer interacts intimately with nine neighbouring gp49 major capsid subunits (Fig. 2, Extended Data Fig. 2e). The N-terminal arm of each gp48 decoration protein sits within a hole formed between the axial domain and extended loop of two neighbouring gp49 proteins (Extended Data Fig. 2f). The flexibility of the N-terminal arm is able to accommodate conformational differences in capsid hexameric faces and pentameric vertices.

The **portal** adopts a unique position within the head and breaks the perfect icosahedral capsid symmetry (Fig. 2b, Extended Data Fig. 2j). Experimentally, it was located within the capsid by 3D classification (see methods) and was reconstructed to 2.6 Å resolution with C12 symmetry imposed (Fig. 2b, c). Each gp45 portal protein monomer adopts the typical bacteriophage portal fold[29], consisting of wing (residues 38-215), clip (residues 253-309), stem (residues 228-253 and 310-331), crown (residues 400-469), and channel valve domains (residues 332-366) (Fig. 1c, 2c, Extended Data Fig. 2g). The portal complex has a minimum inner aperture diameter of 28 Å, which is sufficient to allow DNA to pass through without a conformational change in the portal protein (Extended Data Fig. 2h).

There is a symmetry mismatch where the dodecameric portal interfaces with the surrounding pentameric capsid vertex (Extended Data Fig. 2j). The asymmetric reconstruction of the portal shows that the gp48 capsid decoration protein trimers surrounding the portal are still complete trimers, although density for the 15 residue N-terminal tail is not resolved (Extended Data Fig. 2k). This is reminiscent of the arrangement in Pam3[30]. Presumably the relatively large number of portal proteins (12) allows for subtle conformational plasticity in the symmetry transition from the fivefold capsid opening to the sixfold neck and tail. The portal interacts with the capsid *via* charge-charge interactions between flexible loops made up of residues 164-172 within the gp49 capsomer and 190-193 in the wing motif of the gp45 portal. Interactions are also formed between the stem of the gp45 portal and the gp48 capsid decoration protein.

***Structure of neck reveals a novel portal interacting protein, as well as some similarity to diverse CISs***

The first complex in the neck is the **portal adaptor** gp50, which acts to resolve a near-perfect twelvefold to sixfold symmetry mismatch between the portal and tail (Fig. 1, 2b, Extended Data Fig. 3a). We are not aware that gp50 has any structural homology to other known phage neck protein structures (as determined by Dali, Extended Data Table 3); it is formed of an α-helical bundle, a short β-hairpin, and a C-terminal extension of 11 residues (Extended Data Fig. 3a). The C-terminal extension of gp50 binds directly to the portal, and is tightly pinched between two portal clip domains, forming an extended 5-stranded β-sheet (Fig. 2c, Extended Data Fig. 3b, c). Overall, the interaction between each gp50 portal adaptor C-terminal extension and the three portal chains has a total buried surface of ~1200 $Å^2$, ensuring that the capsid remains attached to the neck. The β-hairpin of gp50 (residues 62 to 71) forms a β-barrel in the dodecameric assembly, and accommodates a decrease in symmetry from C12 to C6 between the gp50 and gp51 (Extended Data Fig. 3e, f). The outside surface of the barrel contains two tyrosine residues per gp50 monomer ($Tyr^{63}$ and $Tyr^{70}$), which face the inner lining of the gp51 **neck valve** hexameric ring (Extended Data Fig. 3e, f). The interacting surface of gp51 is lined with hydrophobic residues that interact with the tyrosines to form a hydrophobic interface (Extended Data Fig. 3e, f). The interface also contains a salt bridge between $Glu^{68}$ from two neighbouring portal adaptor (gp50) chains and arginine residues from each neck valve (gp51) chain, $Arg^{7}$, and $Arg^{48}$ (Extended Data Fig. 3f).

The **neck valve protein** gp51 forms a hexameric complex and forms the narrowest constriction in the neck lumen, ~23 Å in diameter (Extended Data Fig. 3d, e). gp51 has structural homology to the stopper protein of the *Rhodobacter capsulatus* gene transfer agent complex[31], and is made up of an N-terminal helix followed by a β-sheet which bends to form a saddle so that each gp51 monomer sits over three gp53 tail adaptor monomers (Extended Data Fig. 3d).

The **neck-tail adaptor** gp53 forms a hexameric ring linking the neck to the tail (Fig. 2c, Extended Data Fig. 3g, h), and has two tandem domains. Each domain has a similar fold (RMSD 3.2 Å), with an additional C-terminal extension in the tail-proximal domain II. The C-terminus contains a linker and a β-strand, which inserts into the first layer of the tail sheath (Extended Data Fig. 3h, i). The linker between domain II and the C-terminal β-strand is formed of 4 residues (Extended Data Fig. 3c), compared with the 10-13 seen in recently determined pyocin structures[32], AFP[33], and PVC[34] complexes. The β-strand sequence is conserved between the neck-tail adaptor and the tail sheath protein gp55, suggesting that the sequence is important for forming the extended interaction between this region with the sheath domain I β-sheet (Extended Data Fig. 3j).

In order to determine if any large conformational changes are required in the neck proteins to release DNA, the structures in the extended and contracted form were compared (Fig 1). The overall RMSD between all proteins of the neck in the extended state compared to the contracted state is 0.34 Å, indicating that no large conformational change occurs in this region during phage contraction.

***The structure and helical arrangement of the tail subunits is similar to typical CISs in the extended state***

The main **tail tube protein**, gp56, consists of a 7-strand β-sandwich with the conserved fold seen in other *Caudovirales* tail tubes[35-38] (Fig. 1, Extended Data Fig. 4a, b). A loop formed of residues 39 to 59 connects each tail tube ring to the ring below, and a short N-terminal helix of one subunit sandwiches each loop of the subunit above, from the outside (Extended Data Fig. 4a, b). The loop of each tail tube protein interacts with the two neighbouring proteins in the same ring plane, and three proteins in the next ring down, forming extensive interactions as seen in other tailed phages (Extended Data Fig. 4b, c). However, the gp56 tail tube protein lacks both the α-loop and N-loop seen in other myovirus tail tube proteins, as well as the C-terminal arm seen in siphovirus tail tube proteins[38] (Extended Data Fig. 4d). This results in a more open packing between rings of the tail tube (Extended Data Fig. 4c). This is also reflected in the PISA analysis of buried surface between tail tube monomers, with a reduced buried surface of ~1900 Å$^2$ between rings, compared with ~2600 Å$^2$ for AFP, ~2500 Å$^2$ for pyocin, and ~3000 Å$^2$ for T4[32,39,40].

The gp55 **tail sheath protein** is made up of three domains (Fig. 3c). The tail tube proximal C-terminal domain I contains 2 α-helices and a β-sheet. The β-sheet is augmented by N-terminal and C-terminal β-strand insertions from neighbouring sheath proteins, forming an interwoven mesh network (Fig. 3b)[41]. Domain I also contains a loop ('X-loop') consisting of residues 368 to 378 (magenta), which is longer in φCD508 than in the sheath domains of other CISs studied (6 residues and 3 residues respectively) (Extended Data Fig. 4e). This loop pinches the linker of the N-terminal β-strand insertion of a neighbouring sheath protein, reducing the possible range of motion of the gp55 sheath subunits. Domain II contains a β-sheet surrounded by short helices, and forms interactions with neighbouring sheath domains I and II through charge interactions. Domain III is made up of a β-sandwich with an α-helix insertion and appears less ordered than domain I and II in the extended tail reconstructions as evidenced by the decrease in resolution of this region (Extended Data Fig. 5a, b). However, domain III could be fitted from the Rosetta predicted structure placed in the density for the first sheath ring next to the baseplate, in which stabilising interactions form between the sheath distal domain III and the baseplate (Extended Data Fig. 5c).

***φCD508 contains a minimal phage baseplate***

The baseplate of φCD508 is the most complex part of the phage, despite having significantly fewer components than T4 phage[42] (Fig. 2e). In the extended tail conformation, the baseplate forms a hexagonal assembly with a **hub complex** of tail initiator proteins (gp61, gp64) (Extended Data Fig. 6a) towards the centre, a **wedge complex** (gp65a, gp65b, and gp66) (Extended Data Fig. 6c) and **peripheral baseplate** (gp67 and gp68, not modelled) components attached radially towards the outside, with a disk-like side profile (Extended Data Fig. 6c).

The hub complex acts to bind the tail tube and sheath to the baseplate, and the organisation of hub proteins is conserved between φCD508 and other CISs[32-34,42]. One component of the complex is the **tail tube initiator protein** gp61. This protein binds to the subunits in the first layer of the tail tube (gp51), with which it shares a similar β-sandwich fold (Extended Data Fig. 6a). At the C-terminal end of the β-sandwich fold in gp61, a loop (residues 138-165) links to a LysM domain (residues 165-222; Extended Data Fig. 6a, b). The fusion of the LysM domain with the tail tube initiator is similar to that of AFP and PVC[33,34], but different to T4 in which the LysM domain and tail tube initiator are separate proteins.

A second protein in the hub complex is the **sheath initiator protein** gp64. This binds to the terminal ring of sheath proteins and has a similar fold to the sheath protein (gp56) domain I (Extended Data Fig. 6a). A β-sheet made up of 2 β-strands from gp64, binds β-strands donated from two nearby terminal sheath proteins, gp56 (Extended Data Fig. 6a). The C-terminal tail forms contacts with the gp61 tail tube initiator's loop between its β-sandwich domain and lysM domain, stabilising the tail tube and sheath initiation complex (Extended Data Fig. 6b).

The baseplate hub complex also provides a platform onto which the **wedge** is assembled (Fig. 2e, Extended Data Fig. 6b). The wedge consists of three proteins, two gp65 molecules called triplex 1a and triplex 1b, and one gp66 molecule called triplex 2 (Extended Data Fig. 6c, d). The wedge proteins are involved in triggering the contraction of the baseplate, and in the tail-extended state bind to each other component of the baseplate, as well as the needle and tail. Each protein comes together via a core bundle (residues 1-63 and 1-47 for gp65 and gp66 respectively) to form the triplex assembly. The core bundle binds to the hub complex, and is made up of a triple stranded parallel α-helical coiled-coil with a typical core of leucine and isoleucine residues[43].

gp65 has a 'wing' domain (residues 80-184), absent in gp66 (Extended Data Fig. 6e). The wing domain is structurally related to the T4 gp6 wing domain[42], and in the tail-extended state, the respective gp65a and gp65b wing domains adopt a different conformation to bind the tail and needle respectively (Extended Data Fig. 6d). In order to bind two different proteins, flexible loops in the wing domains can alter their conformation (Extended Data Fig. 6e). In triplex 1a, the wing contacts the distal domain III of the first sheath protein in the tail. In triplex 1b, alternating wing domains bind to two different sites on the needle hub and tip, breaking the C6 symmetry of the baseplate and resolving with the C3 symmetry of the needle (Extended Data Fig. 6i).

Both gp65 and gp66 have trifurcation units (residues 64-79 and 185-281, and 47-156 for gp65 and gp66 respectively). These units form a splayed three-pointed star at the end of the core bundle, allowing each protein's C-terminus to point non-symmetrically in different directions (Extended Data Fig. 6d). The C-termini of the two gp65 monomers form dimerisation domains, and six pairs of gp65 dimerisation subunits assemble to form an iris-like circular complex (Extended Data Fig. 6c). The C-terminus of gp66 is predicted to bind the tail fibre gp67, although this interaction is not resolved in our reconstructions (Extended Data Fig. 6f).

***The φCD508 needle complex lacks enzymatic domains, and the typical β-helix observed in other CIS needles***

The **phage needle** is formed of two proteins, a positively charged **needle** protein (gp62), and a pointed **needle tip** (gp63) (Fig. 2e, Extended Data Fig. 6h, 7). The needle protein gp62 functions to resolve the symmetry mismatch between the C6 tail and the C3 needle, forming a hollow trimeric collar around the helical bundle domain of the tip protein (Extended Data Fig. 6h).

The structure of needle protein gp62 is related to a number of other needle proteins, including gp27 of T4 phage[42] (Extended Data Table 2). Each gp62 hub protein forms 2 β-sandwiches, providing six pseudo-symmetric domains to bind to the sixfold symmetric tail tube initiator complex (Extended Data Fig. 6h, 7). Each β-sandwich also contains a lobe (residues 89 to 179 and 196 to 261) which forms a collar surrounding the needle tip (Extended Data Fig. 6h, 7b). gp62 lacks any predicted enzymatic domains such as the lysozyme or peptidoglycan hydrolase domains observed in other CISs[44,45].

The trimeric needle tip protein gp63 (Extended Data Fig. 6h, 7a) adopts a VGR-like fold[46] that differs from that in other phages[47]; it consists of an N-terminal helix (residues 6-25), an oligonucleotide/ oligosaccharide binding (OB-fold) domain (residues 28-55 and 96-114), and an apex domain (residues 56-95). gp63 lacks the β-helix present in other CIS needles[32-34,42,47], and instead the OB-fold attaches directly to the apex domain (Extended Data Fig. 6h, 9a). The apex domain is formed of a short β-hairpin made up of 18 residues, with a HxH motif pointing inwards ($His^{78}$ and $His^{80}$ from each chain) (Extended Data Fig. 6h, 7a). This HxH motif is also found across many CIS needles, and has been shown to bind an iron in other needle tips with similar arrangements of His pairs[47]. A strong density feature is also present between $His^{78}$ and $His^{80}$ residues in gp63, that are predicted to represent a metal binding site with a coordinated ion which we have modelled as an iron ion (Extended Data Fig. 7a).

The N-terminus of gp63 forms a 19 residue helix which resides within the tail tube (Extended Data Fig. 6h, 7b), forming an internal trimeric helical bundle. Augmenting this bundle are three helices at the C-terminus of the tape measure protein, gp59. There is clear interpretable density for 23 residues within the lumen of the tail tube, the remainder of the tape measure protein appearing disordered. Thus, the six bundled helices form a link between the needle tip and the tape measure protein, with the needle tip acting as a plug prior to contraction and genome release (Extended Data Fig. 7).

We have identified >300 protein homologues to the needle tip protein gp63 in putative *Clostridioides* prophage genome sequences and 6 previously characterised *C. difficile* phages based on sequence similarity, showing that the reduced needle tip is widespread among phages of the genus (Extended Data Fig. 9b). We also found that, although low in sequence similarity, the diffocin protein RtbH is also predicted to form a compact needle (Extended Data Fig. 9c)[48]. Further, we were able to identify a similar compact needle tip in phage 3-SAC12 (Extended Data Fig. 9c), which also has a predicted reduced contraction ratio, and infects *L. brevis and other Lactobacilli* [49]. Therefore, this more compact form of the needle may be an adaptation in phage infecting S-layer producing bacteria, although the advantage of this adaptation remains to be determined.

***Tape measure protein***

The putative tape measure protein of φCD508 (gp59) contains multiple hydrophobic and predicted transmembrane stretches as observed in other CIS tape measure proteins[50,51] (Extended Data Fig. 8). The C-terminus of the tape measure protein is predicted to form a number of short β-strands followed by an α-helical 'lazo' domain, which could be modelled in the C3 baseplate/needle reconstruction (Extended Data Fig. 7b, c). As phage tape measure proteins have been associated with multiple functions[52,53], it may be that an increased tail length allows for more functions to be encoded in the tape with implications for infection of these phage. It is intriguing to note that *C. difficile*-infecting phages come in distinct lengths[54,55], including short-tailed (~100 nm), medium-tailed (~130 nm), and long-tailed (>150 nm), and so these classes may encode different functions within their tape measure proteins.

***The contraction ratio of φCD508 bound to receptor is much reduced compared to other CISs***

In order to confirm that reduced contraction of φCD508 in the presence of urea represents the fully contracted state, phage were spontaneously contracted in solution by storing particles for extended periods at 4 °C, thermally contracted by heating to 70 °C, and naturally contracted by binding to fragments of S-layer (Fig. 5). In all cases, TEM showed that phage contracted by 20%.

To determine how φCD508 contracts when binding to the phage's natural host receptor, the S-layer[56], tomograms were collected from samples frozen 40 minutes (n=25) after mixing phage with S-layer fragments (Fig. 5). The lengths of extended and contracted phage tails were measured, and the averages were in agreement with the high resolution structures of φCD508. The extended and contracted tail lengths were approximately 217 nm, and 180 nm respectively (Fig. 1).

The presence or absence of DNA in the phage capsid was also determined. After 5 minutes, 36% of capsids were full (n=239), ~5% were partially empty (n=34), and ~59% were empty (n=395), whereas after 40 minutes ~9% were full (n=49), ~3% were partially empty (n=17), and ~88% were empty (n=481). These results demonstrate that phage contraction is not immediately followed by genome release and suggest that there are additional step(s) between contraction of the sheath and release of DNA from the capsid.

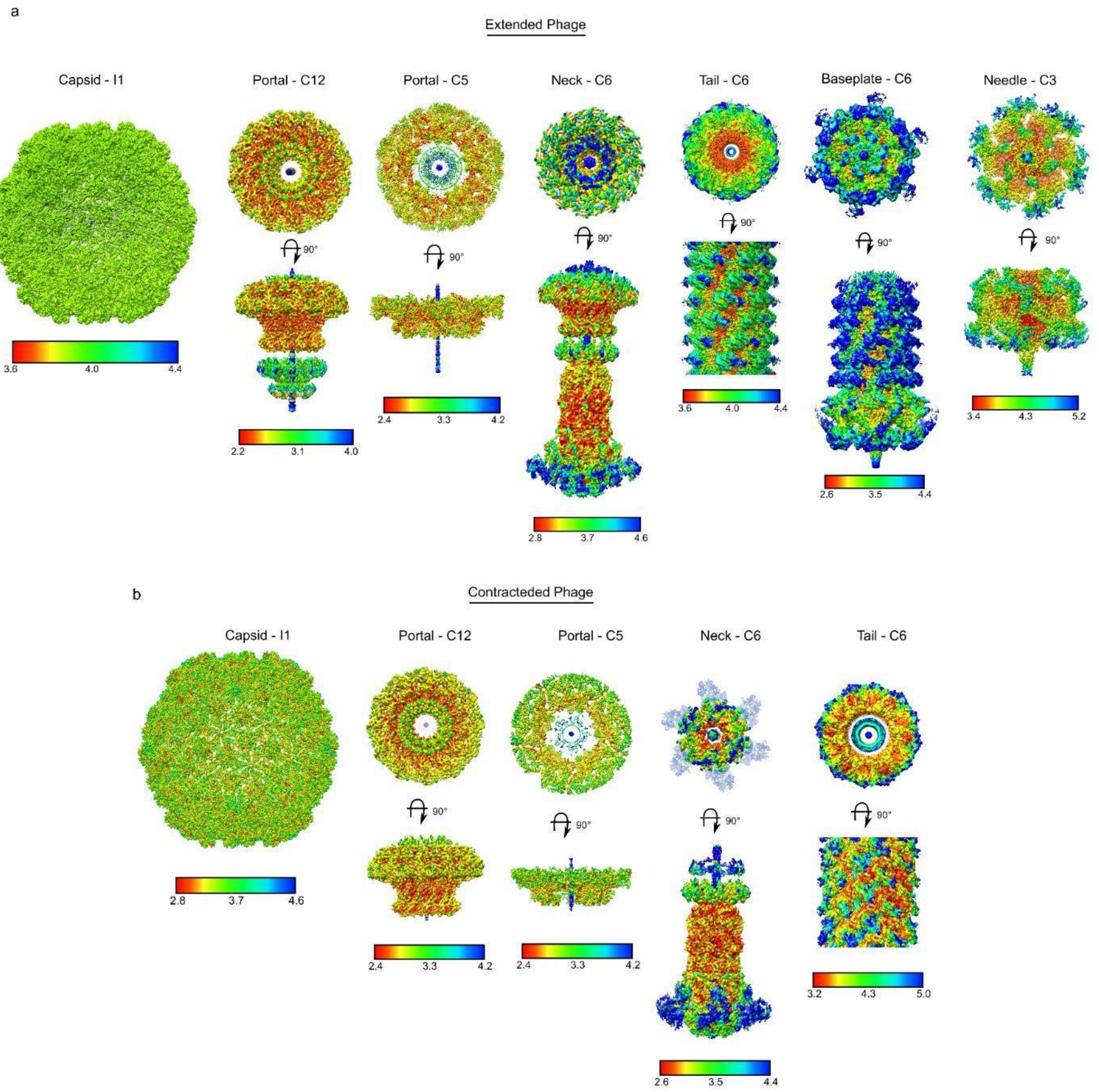


**Extended Data Figure 1 | CryoEM map local resolution plots. a.** CryoEM reconstruction maps for extended φCD508 phage, coloured based on the resolution estimate at each voxel. Plots were calculated using Cryosparc, and coloured from low resolution (blue) to high resolution (red) with scales given for each map.

**b.** CryoEM reconstruction maps for contracted φCD508 phage, coloured based on the resolution estimate at each voxel. Plots were calculated using Cryosparc, and coloured from low resolution (blue) to high resolution (red) with scales given for each map.

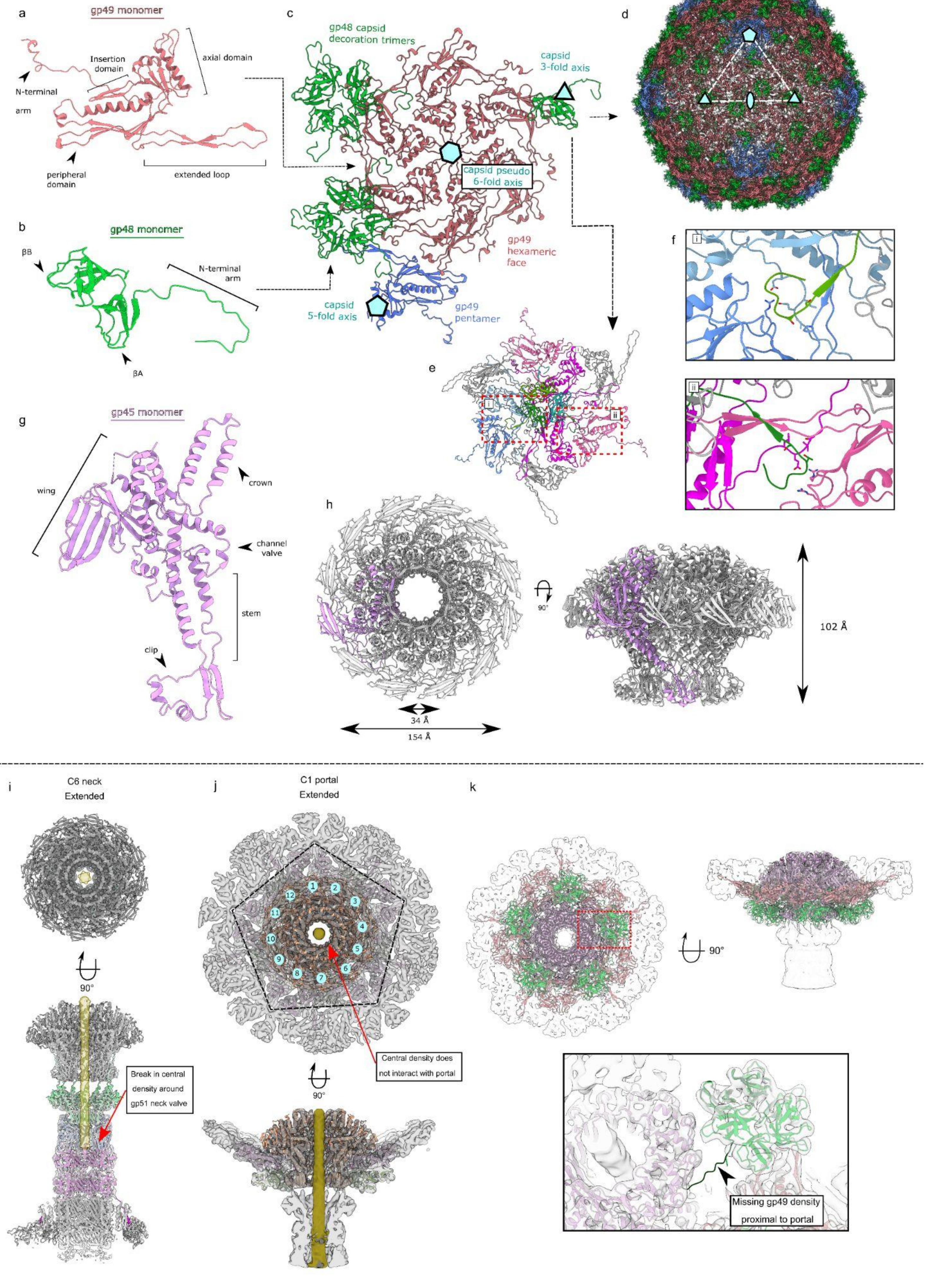
a
gp49 monomer
N-terminal arm
Insertion domain
axial domain
peripheral domain
extended loop
b
gp48 monomer
βB
N-terminal arm
βA
c
gp48 capsid decoration trimers
capsid 3-fold axis
capsid pseudo 6-fold axis
gp49 hexameric face
capsid 5-fold axis
gp49 pentamer
d
e
f
g
gp45 monomer
wing
crown
channel valve
stem
clip
h
90°
102 Å
34 Å
154 Å
i
C6 neck Extended
90°
Break in central density around gp51 neck valve
j
C1 portal Extended
Central density does not interact with portal
90°
k
90°
Missing gp49 density proximal to portal

**Extended Data Figure 2 | Capsid and portal interactions**. **a.** Cartoon representation of gp49 major capsid protein with domain features highlighted. **b.** Cartoon representation of gp48 capsid decoration protein monomer with domain features highlighted. **c.** cartoon representation of capsomer asymmetric unit with gp49 penton vertex protein in blue. Symmetry and pseudosymmetry axes are highlighted. **d.** Cartoon representation of assembled capsid, coloured as in a-c. **e.** The organisation of nine gp49 capsid proteins that interact with each gp48 trimer. **f.** Insets show specific interactions formed between the capsid decoration and major capsid proteins at the (i) pentamer vertex and (ii) hexamer face. **g.** Cartoon representation of the gp45 portal protein monomer with domain features highlighted. **h.** Portal dodecameric assembly with portal dimensions highlighted. **i.** CryoEM density of C6 neck reconstruction with cartoon models also shown. DNA density is highlighted in yellow shown to stop at the gp51 neck valve assembly. **j.** CryoEM density of C1 portal/capsid assembly showing the symmetry mismatch between the pentamer capsid vertex and dodecameric portal. DNA density shown in yellow does not form interactions with any portal proteins. **k.** CryoEM density of C1 portal/capsid assembly showing the symmetry mismatch between gp48 capsid decoration trimers and the dodecameric portal assembly. Inset shows lacking density for the N-terminal arm of one portal proximal gp48 protein.

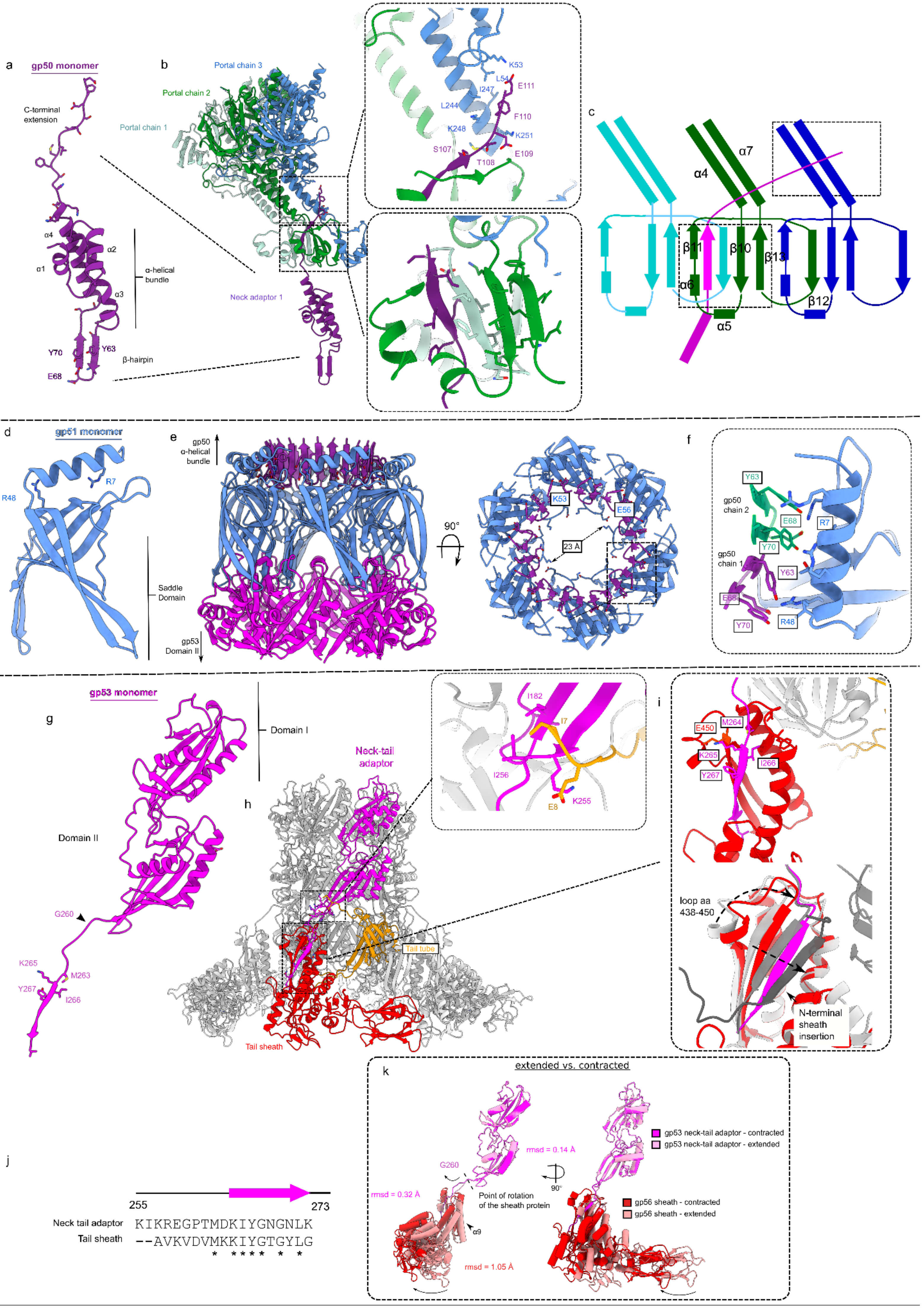

a
gp50 monomer
C-terminal extension
α4
α1
α2
α3
α-helical bundle
Y70
Y63
E68
β-hairpin
b
Portal chain 3
Portal chain 2
Portal chain 1
Neck adaptor 1
K53
L54
I247
E111
L244
F110
K248
K251
S107
T108
E109
c
α7
α4
β11
β10
β13
α6
β12
α5
d
gp51 monomer
R7
R48
Saddle Domain
e
gp50 α-helical bundle
gp53 Domain II
90°
K53
E56
23 Å
f
Y63
gp50 chain 2
E68
R7
Y70
gp50 chain 1
Y63
E68
Y70
R48
g
gp53 monomer
Domain I
Domain II
G260
K265
M263
Y267
I266
h
Neck-tail adaptor
Tail tube
Tail sheath
I182
I7
I256
E8
K255
i
E450
M264
K265
I266
Y267
loop aa 438-450
N-terminal sheath insertion
j
255
273
Neck tail adaptor KIKREGPTMDKIYGNGNLK
Tail sheath --AVKVDVMKKIYGTGYLG
* **** * *
k
extended vs. contracted
rmsd = 0.14 Å
G260
rmsd = 0.32 Å
Point of rotation of the sheath protein
α9
rmsd = 1.05 Å
90°
gp53 neck-tail adaptor - contracted
gp53 neck-tail adaptor - extended
gp56 sheath - contracted
gp56 sheath - extended

**Extended Data Figure 3 | Neck interactions**. **a.** Cartoon representation of gp50 portal adaptor protein monomer with domain features highlighted. **b.** Cartoon representation of gp51 monomer interacting with three unique portal chains. Insets show specific interactions between (i) C-terminal extension of gp50 and portal chain 2 and 3 stem domains, and (ii) β-sheet formed by insertion of C-terminal extension of gp50 into portal chain 1 and 2 clip domains **c.** Schematic representation of interactions highlighted in b. **d.** Cartoon representation of gp51 neck valve protein monomer with domain features highlighted. **e.** Cartoon representation of hexameric assembly of gp51, and interactions with gp50 (purple) and gp53 (magenta) **f.** Inset shows symmetry mismatch interactions between dodecameric gp50 and hexameric gp51. **g.** Cartoon representation of the gp53 portal protein monomer with domain features highlighted. **h.** Cartoon representation of hexameric assembly of gp53, and interactions formed with sheath protein (red), and tail tube protein (orange). Inset shows specific interaction between gp53 and tail tube protein. **i.** Comparison of gp53 β-strand interaction with the terminal sheath protein (top) with intermediate sheath protein mesh network (bottom, grey). **j.** Sequence similarity between Neck-tail adaptor C-terminal -strand and sheath protein β-strand. **k.** Comparison between the extended (pink) and contracted (red) state of the neck proximal sheath proteins, showing that the movement is a rigid body pivot around Gly260.

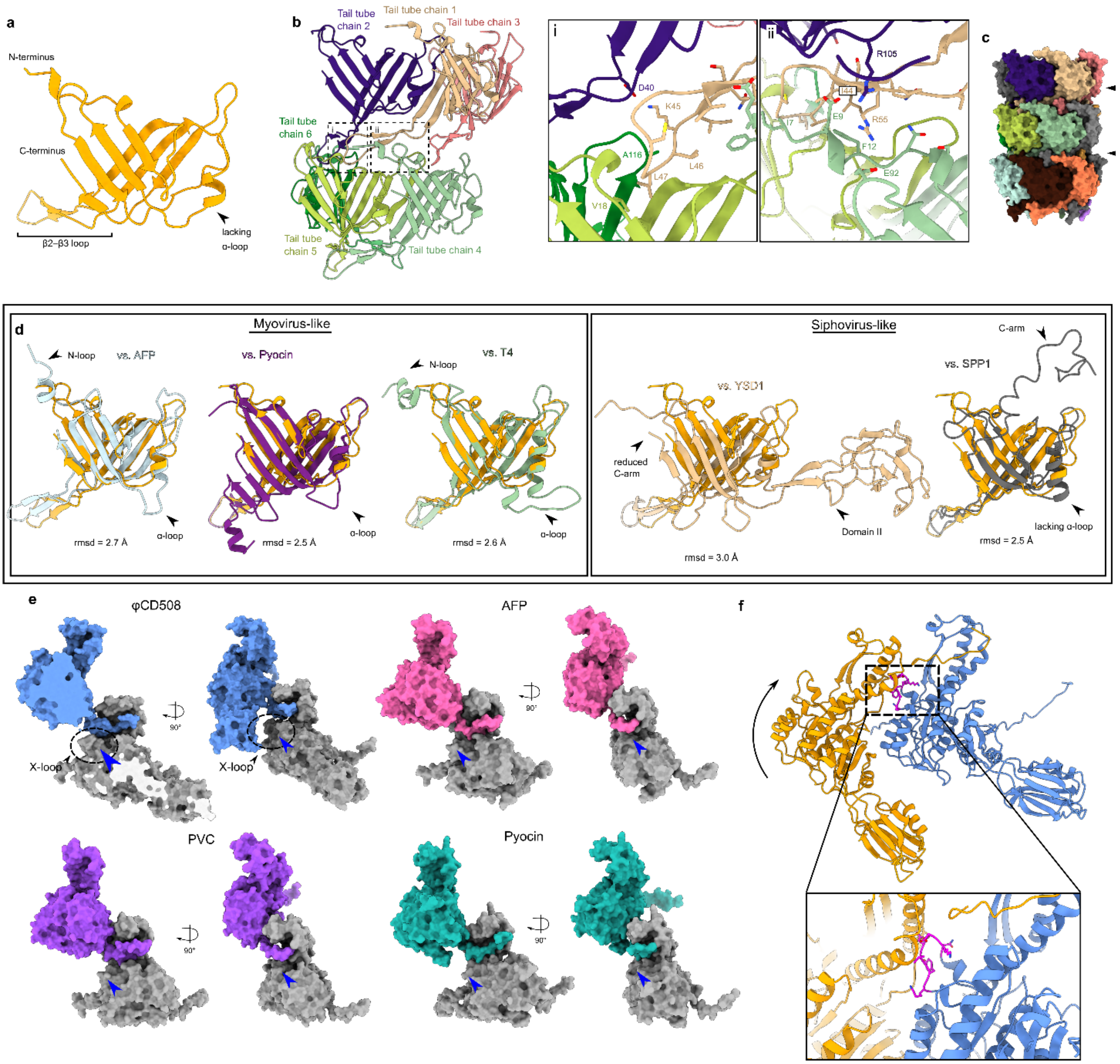


**Extended Data Figure 4 | Sheath and tail tube interactions**. **a.** Cartoon representation of tail tube protein with domain features highlighted. **b.** Cartoon representation of extensive interactions formed between each tail tube protein. Insets show (i) interactions between chain 1 and chains 2, 5, and 6, and (ii) the clamp between chains 2 and 4. **c.** Surface rendering of three layers of tail tube protein, showing the large ridges formed by the lack of an α-loop in gp56 tail tube. **d.** Comparison between φCD508 tail tube protein and known structures for myovirus tail tubes (left), and siphovirus tail tubes (right), with features such as N-loop, α-loop, and C-arm highlighted. **e.** Surface rendering of the sheath N-terminal linker handshake between neighbouring sheath proteins for φCD508 (blue) and myovirus structural homologues. φCD508 contains an X-loop formed of residues 368 to 378, missing in other knownsheath protein structures (blue arrow). **f.** Cartoon model of two sheath proteins as coloured in Figure 3a fitted onto the contracted structure of pyocin sheath proteins. For the conventional contraction extent of 50%, the X-loop (magenta) clashes severely with the neighbouring sheath chain.

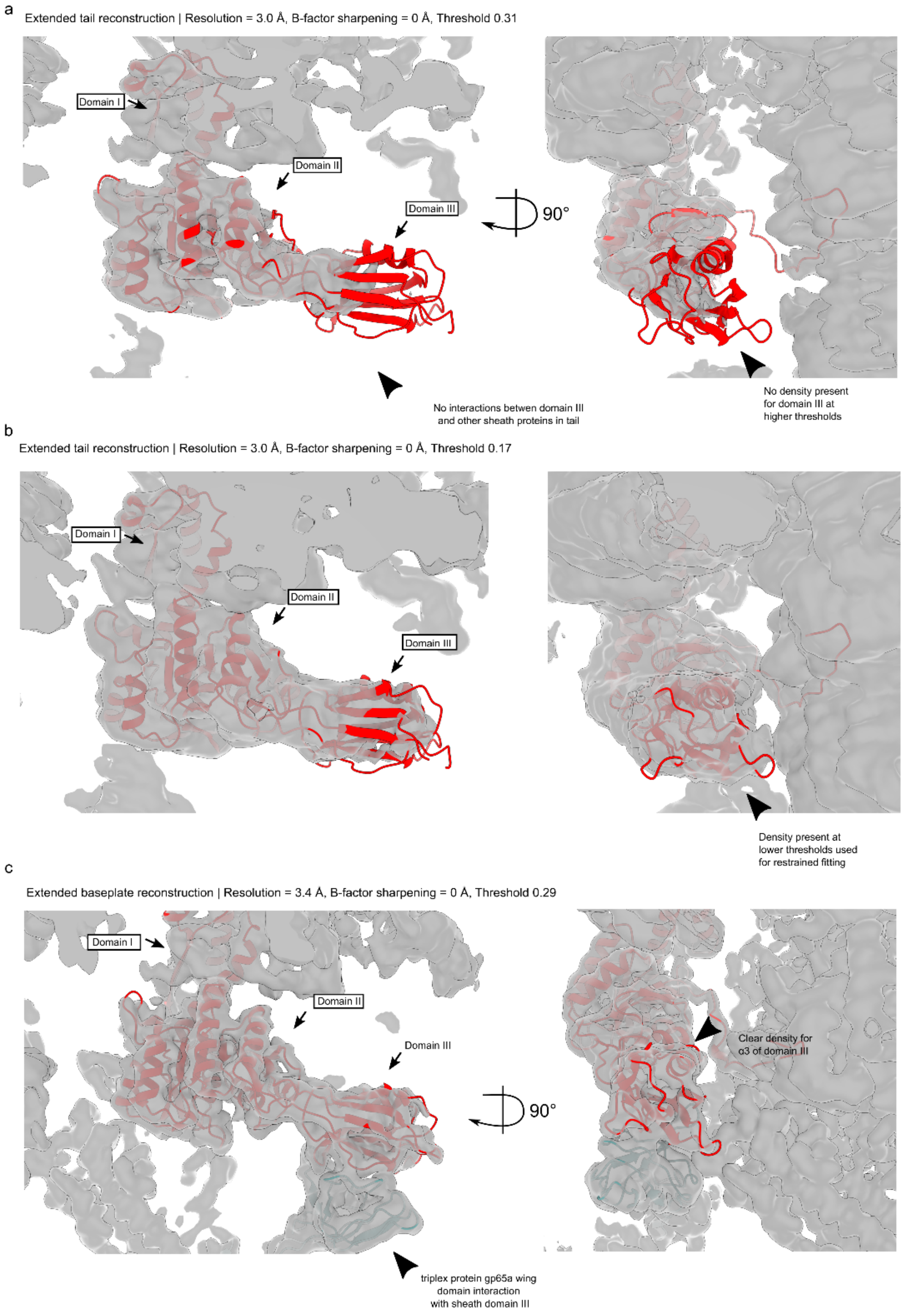


**Extended Data Figure 5 | Sheath CryoEM map density fits.** **a.** Unsharpened sheath protein CryoEM density for the tail reconstruction, thresholded at **σ**=0.31. Domain I and II reside in strong density, but domain III lacks density due to flexibility in the region. **b.** Unsharpened sheath protein CryoEM density for the tail reconstruction, thresholded at **σ**=0.17. Some density is apparent for flexible fitting of the baseplate proximal sheath model, but not for *de novo* model building. **c.** Unsharpened sheath protein CryoEM density for the baseplate proximal sheath layer, thresholded at **σ**=0.29. The association of domain III with the triplex protein gp65a wing domain increases the order of the region such that good density is present for domain III.

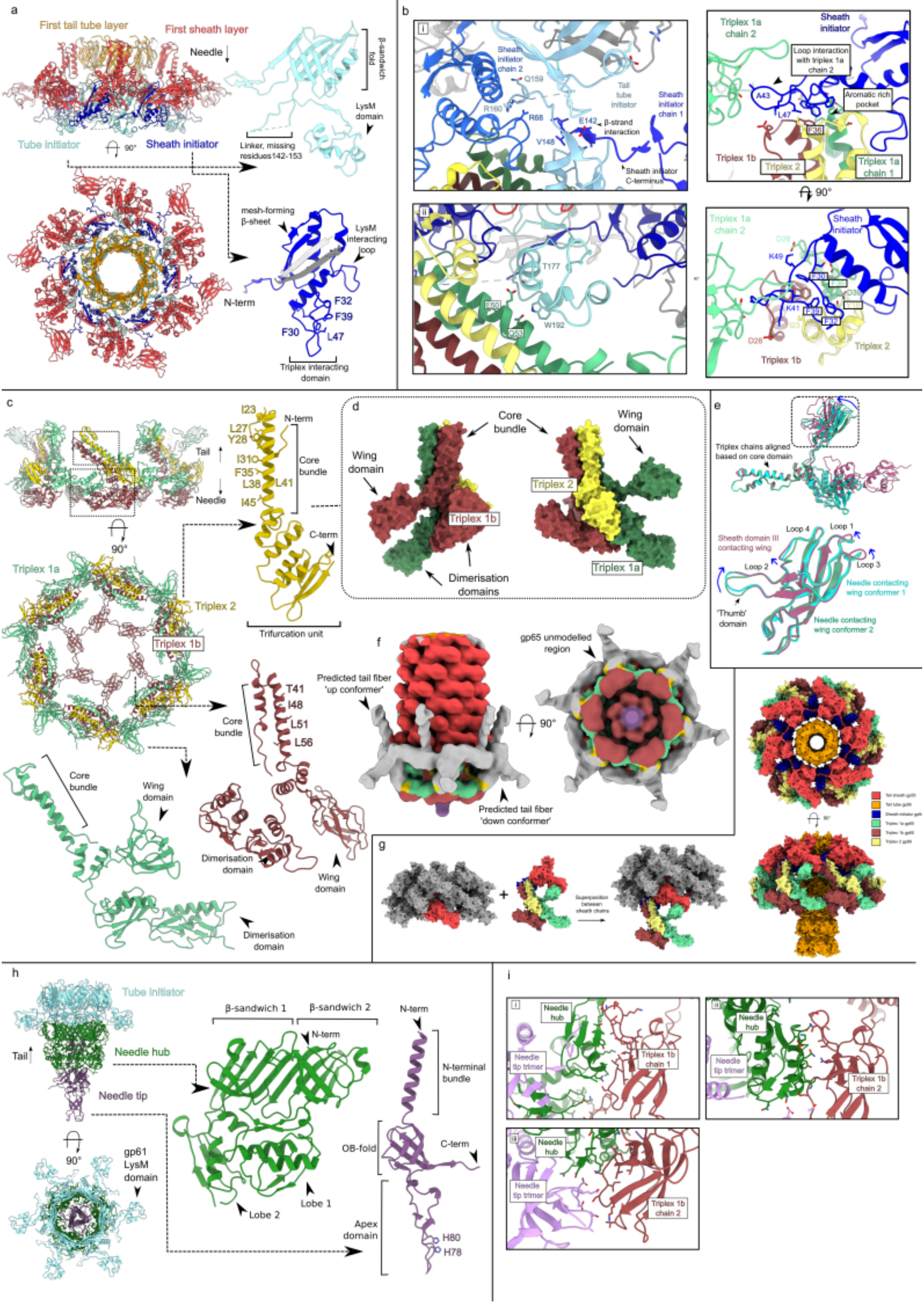

a
First tail tube layer
First sheath layer
Needle
β-sandwich fold
LysM domain
Linker, missing residues142-153
Tube initiator
Sheath initiator
90°
mesh-forming β-sheet
LysM interacting loop
N-term
F32
F39
F30
L47
Triplex interacting domain
b
i
Sheath initiator chain 2
Q159
R160
R68
Tail tube initiator
Sheath initiator chain 1
E142
β-strand interaction
V148
Sheath initiator C-terminus
ii
T177
E60
Q53
W192
Triplex 1a chain 2
Sheath initiator
Loop interaction with triplex 1a chain 2
A43
Aromatic rich pocket
L47
F38
Triplex 1b
Triplex 2
Triplex 1a chain 1
90°
D28
K49
K41
F39
F32
D38
I23
Triplex 1b
Triplex 2
c
Tail
Needle
90°
Triplex 1a
Triplex 2
Triplex 1b
I23
L27
Y28
I31
F35
L38
L41
I45
N-term
Core bundle
C-term
Trifurcation unit
T41
I48
L51
L56
Core bundle
Wing domain
Dimerisation domain
Wing domain
Dimerisation domain
d
Core bundle
Wing domain
Wing domain
Triplex 1b
Triplex 2
Triplex 1a
Dimerisation domains
e
Triplex chains aligned based on core domain
Loop 4
Loop 1
Sheath domain III contacting wing
Loop 2
Loop 3
Needle contacting wing conformer 1
'Thumb' domain
Needle contacting wing conformer 2
f
Predicted tail fiber 'up conformer'
gp65 unmodelled region
90°
Predicted tail fiber 'down conformer'
g
Superposition between sheath chains
90°
h
Tube initiator
Needle hub
Tail
Needle tip
90°
gp61 LysM domain
β-sandwich 1
β-sandwich 2
N-term
Lobe 2
Lobe 1
N-term
N-terminal bundle
OB-fold
C-term
Apex domain
H80
H78
i
Needle hub
Needle tip trimer
Triplex 1b chain 1
Needle hub
Needle tip trimer
Triplex 1b chain 2
Needle hub
Needle tip trimer
Triplex 1b chain 2

**Extended Data Figure 6 | Baseplate interactions a.** Cartoon representation of baseplate hub proteins assembled (left) including the first layer of tail tube protein (orange) and sheath protein (red), and in monomeric representation with domain features highlighted (right). **b.** Insets showing interactions between sheath initiator protein and tail tube initiator protein (top left), and between tail tube initiator and triplex proteins in baseplate wedge (bottom left). Right inset shows how baseplate wedge is assembled on the sheath initiator protein, and how movement of the wedge is propagated into the sheath proteins via the sheath initiator. **c.** Cartoon representation of baseplate wedge proteins assembled (left) and in monomeric representations with domain features highlighted. **d.** Surface rendering of triplex complex showing the three main features. **e.** Conformational variability in the wing domain within three conformers of triplex 1 protein. **f.** Baseplate CryoEM reconstruction lowpass filtered to 20 Å, and thresholded at σ=0.022, showing density for putative tail fibers in an upwards and downwards conformation. **g.** Surface rendered model of φCD508 baseplate contraction based on contracted sheath reconstruction, and extended baseplate reconstruction. Each protein is coloured as in a and c. **h.** Cartoon representation of needle proteins assembled (left), with the tail tube initiator proteins also present (cyan) for reference, and in monomeric representations (right) with domain features highlighted. **i.** Insets show interactions between the triplex 1a wing domains with the needle tip and needle hub proteins, in two forms due to symmetry mismatch between C3 needle and C6 baseplate hub proteins.

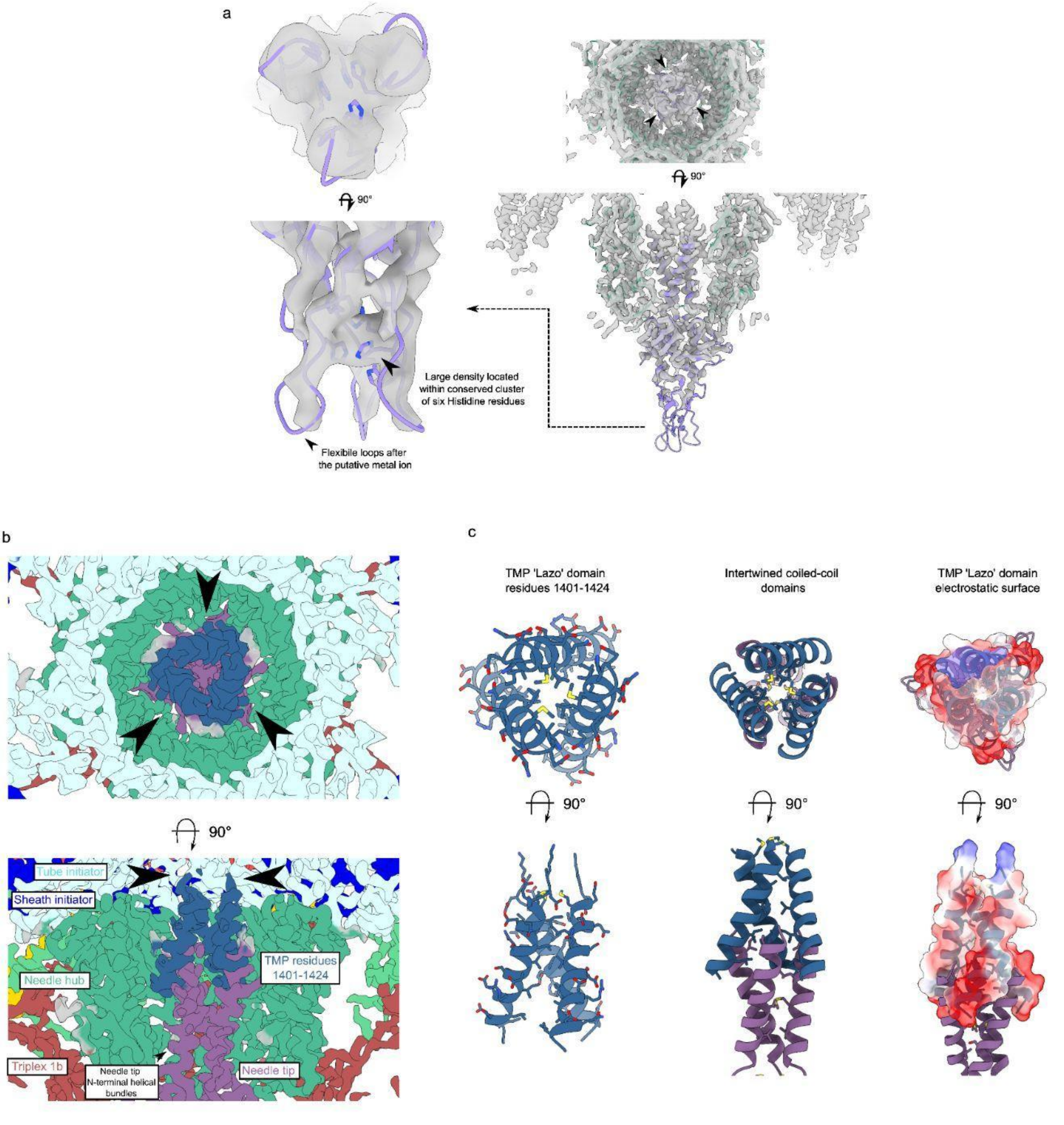


**Extended Data Figure 7 | Needle and tape measure protein interactions a.** CryoEM density map of needle tip apex domain with needle tip trimer (purple) and needle hub (green). Histidine residues surround a strong density feature interpreted as a metal ion. **b.** Slices through CryoEM needle C3 reconstruction coloured based on protein. Clear helical density with sidechain density is visible for the TMP lazo domain (dark blue). **c.** Cartoon representation of tape measure protein trimer with sidechains shown (left). Interaction between tape measure protein and the N-terminal bundle domain of the needle tip. The coiled-coils are formed of three interleaved leucine zippered splayed bundles (middle). Electrostatic potential of tape measure protein lazo domain with strong negative charge present proximal to the needle tip protein.

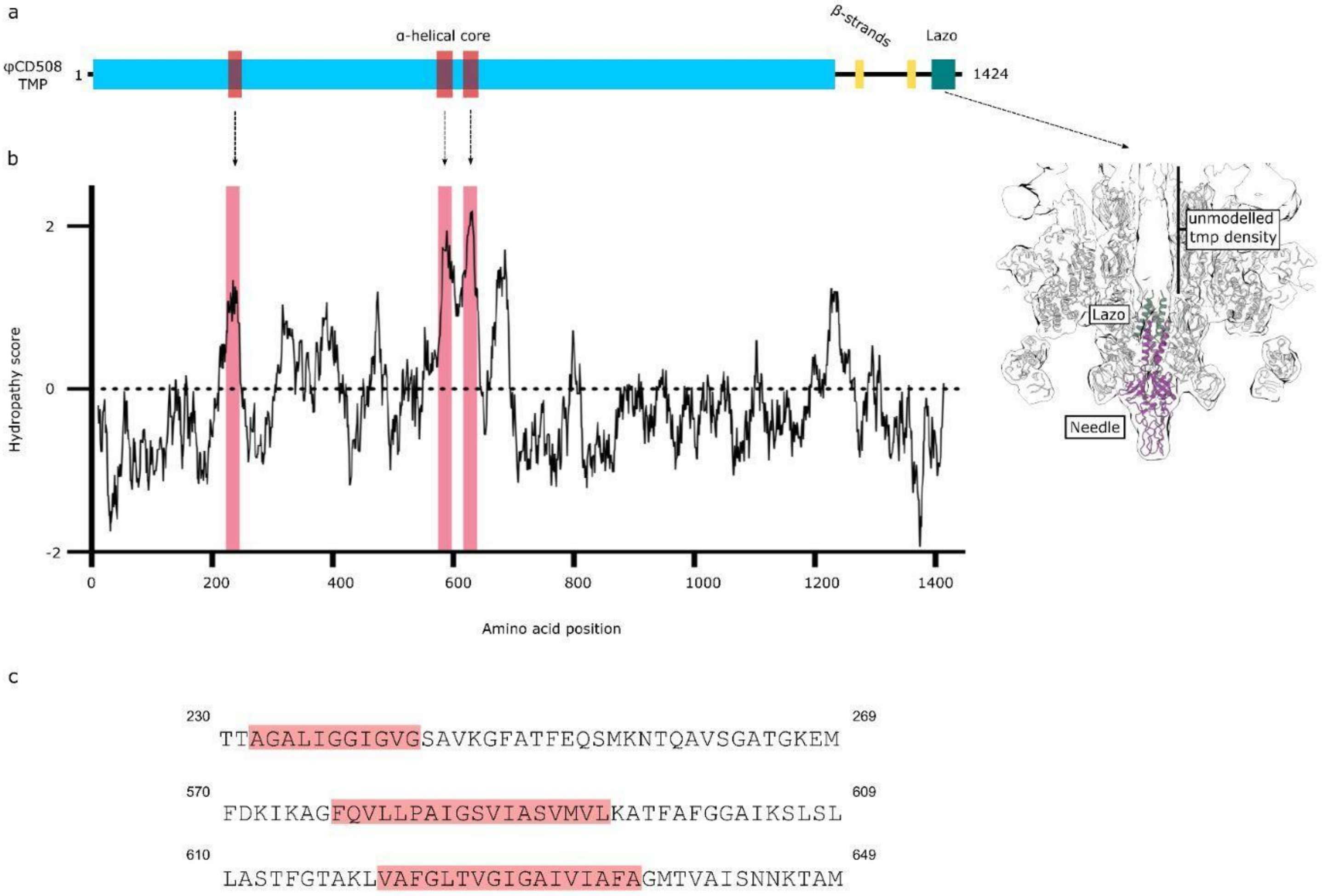


**Extended Data Figure 9 | Tape measure protein features**

**a.** Schematic diagram of φCD508 tape measure protein based on Psipred secondary structure prediction and previous characterisation of tape measure proteins. **b.** Kyte and Doolittle hydropathy analysis[57] of the tape measure protein gp59, performed using Expasy ProtScale[59] with window size 21. More hydrophobic regions of the protein are shown above the dotted horizontal line at 0. The three putative transmembrane helices shown in a., predicted using DeepTMHMM[58], are highlighted in pink**. c.** Amino acid sequence of predicted transmembrane domains with hydrophobic transmembrane stretches highlighted in red.

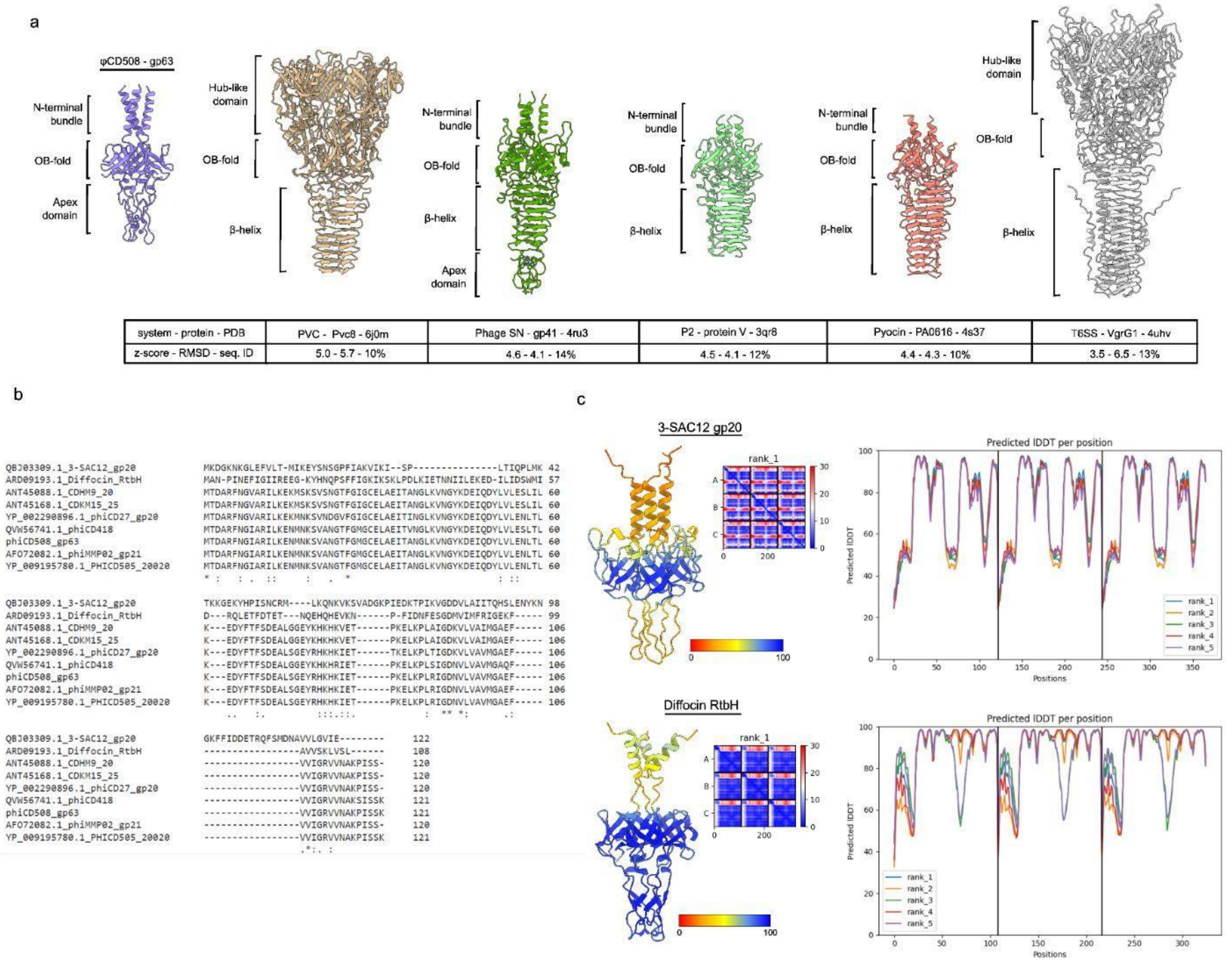


**Extended Data Figure 8 | Needle structural homologues and family features a.** Structural features of φCD508 needle tip protein gp63 compared with structures determined for homologous needle tip proteins from phage and T6SS, as well as z-score, RMSD, and sequence identity. **b.** Sequence alignment between φCD508 needle tip protein gp63 and phage needle proteins from other bacteriophages infecting S-layer-producing bacterial species. **c.** AlphaFold predicted structures of needle tips from *L. bravis* infecting bacteriophage 30SAC12, and *C. difficile* infecting phage-like particle diffocin . Each needle lacks the β-helix, an omission common to bacteriophages infecting S-layer producing species. Predicted structures are coloured based on pLDDT score, and pLDDT is plotted to show the confidence in the predicted structure (right).

### Extended Data Table 1. Cryo-EM data collection, refinement and validation statistics

| | Extended Phage Capsid I1 (EMDB-xxxx) (PDB xxxx) | Extended Phage Portal C5 (EMDB-xxxx) (PDB xxxx) | Extended Phage Portal C12 (EMDB-xxxx) (PDB xxxx) | Extended Phage Neck (EMDB-xxxx) (PDB xxxx) | Extended Phage Tail (EMDB-xxxx) (PDB xxxx) | Extended Phage Baseplate (EMDB-xxxx) (PDB xxxx) | Extended Phage Needle (EMDB-xxxx) (PDB xxxx) | Contracted Phage Capsid I1 (EMDB-xxxx) (PDB xxxx) | Contracted Phage Portal C12 (EMDB-xxxx) (PDB xxxx) | Contracted Phage Neck (EMDB-xxxx) (PDB xxxx) | Contracted Phage Tail (EMDB-xxxx) (PDB xxxx) |
|---|---|---|---|---|---|---|---|---|---|---|---|
| **Data collection and processing** | | | | | | | | | | | |
| Magnification | 81,000 | 81,000 | 81,000 | 81,000 | 81,000 | 81,000 | 81,000 | 81,000 | 81,000 | 81,000 | 81,000 |
| Voltage (kV) | 300 | 300 | 300 | 300 | 300 | 300 | 300 | 300 | 300 | 300 | 300 |
| Electron exposure (e–/Å$^2$) | 42 | 42 | 42 | 42 | 42 | 42 | 42 | 42 | 42 | 42 | 42 |
| Defocus range (μm) | 1.5-3 | 1.5-3 | 1.5-3 | 1.5-3 | 1.5-3 | 1.5-3 | 1.5-3 | 1.5-3 | 1.5-3 | 1.5-3 | 1.5-3 |
| Pixel size (Å) | 1.06 | 1.06 | 1.06 | 1.06 | 1.06 | 1.06 | 1.06 | 1.06 | 1.06 | 1.06 | 1.06 |
| Symmetry imposed | I1 | C5 | C12 | C6 | C6 + helical | C6 | C3 | I1 | C12 | C6 | C6 |
| Initial particle images (no.) | 61,489 | 70,251 | 70,251 | 68,159 | 178,796 | 19,276 | 19,276 | 42,410 | 31,079 | 30,325 | 128,963 |
| Final particle images (no.) | 61,489 | 23,343 | 38,968 | 23,724 | 157,557 | 19,276 | 17,759 | 38,654 | 26,404 | 30,325 | 48,919 |
| Map resolution (Å) | 3.74 | 3.23 | 2.64 | 3.40 | 2.71 | 3.43 | 3.96 | 3.28 | 2.88 | 3.27 | 4.21 |
| FSC threshold | 0.143 | 0.143 | 0.143 | 0.143 | 0.143 | 0.143 | 0.143 | 0.143 | 0.143 | 0.143 | 0.143 |
| **Refinement** | | | | | | | | | | | |
| Initial model used (PDB code) | *de novo* | *de novo* | *de novo* | *de novo* | *de novo* | *de novo* | *de novo* | *de novo* | *de novo* | *de novo* | *de novo* |
| Model resolution (Å) | | 3.6 | 2.8 | 3.6 | 3.0 | 3.8 | 3.7 | 3.5 | 3.2 | 3.5 | 4.3 |
| FSC threshold | 0.5 | 0.5 | 0.5 | 0.5 | 0.5 | 0.5 | 0.5 | 0.5 | 0.5 | 0.5 | 0.5 |
| Model resolution range (Å) | | | | | | | | | | | |
| Map sharpening *B* factor (Å$^2$) | -160.8 | -71.7 | -114.6 | -86.5 | -77.7 | -100.0 | -103.9 | -146.9 | -94.9 | -81.4 | -194.2 |
| Model composition | 169,271 | 72,277 | 85,788 | 193,800 | 167,202 | 172,440 | 192,782 | 45,235 | 85,788 | 193,800 | 130,436 |
| Non-hydrogen atoms | 84,599 | 36,201 | 43,008 | 97,188 | 83,466 | 85,896 | 95,949 | 22,687 | 43,008 | 97,188 | 65,130 |
| Protein residues | 10,730 | 4,595 | 5,340 | 12,252 | 10,620 | 10,914 | 12,183 | 2,877 | 5,340 | 12,252 | 8,280 |
| Ligands | 0 | 0 | 0 | 0 | 0 | 0 | 0 | 0 | 0 | 0 | 0 |
| *B* factors (Å$^2$) | | | | | | | | | | | |
| Protein | 103.72 | 112.4 | 64.4 | 145.44 | 181.98 | 190.82 | 100.47 | 69.30 | 111.0 | 166.0 | 112.28 |
| Ligand | - | - | - | - | - | - | - | - | - | - | - |
| R.m.s. deviations | | | | | | | | | | | |
| Bond lengths (Å) | 0.004 | 0.004 | 0 | 0.004 | 0.006 | 0.004 | 0.004 | 0.004 | 0.004 | 0.004 | 0.004 |
| Bond angles (°) | 0.744 | 0.766 | 0 | 0.629 | 0.605 | 0.658 | 0.640 | 0.664 | 0.622 | 0.636 | 0.694 |
| Validation | | | | | | | | | | | |
| MolProbity score | 1.51 | 1.32 | 1.36 | 1.44 | 1.48 | 1.28 | 1.38 | 1.37 | 0.89 | 1.46 | 1.35 |
| Clashscore | 2.98 | 2.48 | 6.62 | 4.41 | 4.75 | 2.69 | 2.83 | 2.47 | 1.52 | 4.59 | 2.58 |
| Poor rotamers (%) | 0.00 | 0.05 | 0.04 | 0.16 | 0.09 | 0.00 | 0.00 | 0.00 | 0.00 | 0.01 | 0.00 |
| Ramachandran plot | | | | | | | | | | | |
| Favoured (%) | 93.63 | 95.82 | 98.20 | 96.54 | 96.41 | 96.53 | 95.62 | 95.11 | 98.19 | 96.51 | 95.63 |
| Allowed (%) | 6.31 | 4.11 | 1.80 | 3.44 | 3.59 | 3.47 | 4.38 | 4.71 | 1.81 | 3.49 | 4.26 |
| Disallowed (%) | 0.00 | 0.07 | 0.00 | 0.02 | 0.00 | 0.00 | 0.00 | 0.18 | 0.00 | 0.00 | 0.11 |

### Extended Data Table 2 | φCD508 proteins structural homologues

| Phage region | phiCD508 gene product | Phage/PTLP | Protein | PDB | Z-score | RMSD | Sequence Id % |
|---|---|---|---|---|---|---|---|
| Head | gp45 Portal protein | SPP1 | Portal protein | 2jes | 18.5 | 4.4 | 14 |
| | | G20C | Portal protein | 4zjn | 18.4 | 4.6 | 12 |
| | | RcGTA | Rcc01684 | 6toa | 17.2 | 3.9 | 10 |
| | | T7 | Portal protein | 7bou | 15.9 | 5.1 | 10 |
| | | Pam1 | Pam1 portal protein | 7eep | 15.8 | 5.8 | 9 |
| | | T4 | Portal protein gp20 | 3ja7 | 15.3 | 5 | 8 |
| | | P22 | Portal protein | 5gai | 11.2 | 3.8 | 7 |
| | gp49 Major capsid protein | HK97 | Major capsid protein | 1ohg | 14.8 | 4.2 | 10 |
| | | RcGTA | Rcc01687 | 6tsw | 13.2 | 3.7 | 9 |
| | | YSD1 | YSD1_17 | 6xgq | 13.1 | 4.3 | 4 |
| | | T7 | Major capsid protein 10A | 3j7w | 12.8 | 4.2 | 12 |
| | | T4 | Major capsid protein | 5vf3 | 12.8 | 3.2 | 8 |
| | | Pam1 | Major capsid proteins | 7eel | 12.4 | 4.9 | 9 |
| | gp48 Capsid decoration protein | TW1 | Capsid Stabilizing Protein | 5wk1 | 6.4 | 3.1 | 9 |
| | | XM1 | Minor capsid protein | 7kmx | 6.3 | 3.2 | 15 |
| | | phi29 | gp8.5 Capsid fiber protein | 6qyy | 5.7 | 3.3 | 12 |
| | | YSD1 | YSD1_16 | 6xgq | 5.1 | 14.3 | 8 |
| Neck | gp53 Neck-tail adaptor | RcGTA | RCC01690 | 6te9 | 13.2 | 2.8 | 14 |
| | | Lambda | Minor tail protein U | 3fz2 | 12 | 3.3 | 10 |
| | | Pyocin | PA0615 | 6u5j | 11.6 | 9.7 | 11 |
| | | AFP | Afp16 | 6rap | 8.5 | 6.9 | 9 |
| | | T4 | Gp15 | 3j2m | 8.4 | 6.4 | 8 |
| | | SPP1 | Gp17 | 2lfp | 7.8 | 3.5 | 13 |
| | gp50 Neck-portal adaptor | no known structural homologues | | | | | |
| | gp51 Neck valve protein | RcGTA | RCC01688 | 6te9 | 4 | 3.2 | 7 |
| | | PBSX | PBSX protein XkdH | 3f3b | 2.6 | 4.2 | 5 |
| Tail | gp55 Tail sheath | Pyocin | sheath | 3j9q | 30.3 | 3.3 | 12 |
| | | Diffocin | Putative phage XkdK-like protein | 6gkw | 29.6 | 2.5 | 22 |
| | | AFP | AFP3 | 6rao | 26 | 3.9 | 9 |
| | | T4 | Tail sheath protein Gp18 | 3j2m | 23.8 | 3.9 | 16 |
| | | phi812K1-420 | tail sheath protein | 5li2 | 20.3 | 3.7 | 13 |
| | | T6SS | type VI secretion protein | 5mxn | 18.2 | 9.2 | 9 |
| | gp56 Tail tube | AFP | AFP1 | 6rap | 12 | 2.7 | 11 |
| | | SPP1 | Tail tube protein gp17.1 | 6yeg | 9.9 | 2.5 | 12 |
| | | T4 | Tail tube protein gp19 | 5w5f | 9.8 | 2.6 | 11 |
| | | Pyocin | FIIR2 protein | 5w5e | 9.5 | 2.5 | 14 |
| | | YSD1 | YSD1_22 major tail protein | 6xgr | 9.2 | 3 | 6 |
| | | T5 | Tail tube protein | 5ngj | 8.4 | 2.3 | 13 |
| Baseplate | gp65 Triplex 1a/b | PVC | Pvc11 | 6j0n | | | |
| | | Pyocin | PA0618 | 6u5b | | | |
| | | AFP | Afp11 | 6rao | 12.6 | 16.2 | 9 |
| | | T4 | gp6 | 5hx2 | 8.2 | 12 | 16 |
| | | XM1 | gp16 | 7kh1 | | | |
| | gp61 Tail tube initiator | AFP | Afp7 | 6rao | 10.6 | 10.2 | 13 |
| | | P2 | ORF15 | 2wzp | 9 | 6.2 | 10 |
| | | SPP1 | gp 19.1 | 2x8k | 7.9 | 6.7 | 7 |
| | | RcGTA | | 6toa | 7.6 | 3.7 | 9 |
| | | SPP1 | gp17.1 | 6yeg | 7.1 | 3.1 | 11 |
| | | YSD1 | YSD1_22 | 6xgr | 6.9 | 3.2 | 11 |
| | gp64 Sheath initiator | AFP | Afp9 | 6rao | 9.7 | 2.5 | 21 |
| | | T4 | gp25 | 5iw9 | 7.7 | 5.6 | 11 |
| | | T6SS | TssE | 6gj1 | 6.8 | 3.2 | 11 |
| | | RcGTA | Rcc01690 | 6te9 | 4.4 | 3.4 | 5 |
| | gp66 Triplex 2 | T4 | gp6 | 5hx2 | 6.5 | 7.1 | 8 |
| | | AFP | Afp11 | 6rao | 5.9 | 7.6 | 8 |
| Needle | gp63 Needle tip | PVC | Pvc8 | 6j0m | 5.0 | 5.7 | 10 |
| | | Phage SN | puncturing protein gp41 | 4ru3 | 4.6 | 4.1 | 14 |
| | | P2 | Baseplate assembly protein V | 3qr8 | 4.5 | 4.1 | 12 |
| | | Pyocin | PA0616 | 4s37 | 4.4 | 4.3 | 10 |
| | | T6SS | VgrG1 | 4uhv | 3.5 | 6.5 | 13 |
| | gp62 Needle | Pyocin | hub | 6u5h | 19.6 | 3.0 | 10 |
| | | PVC | Pvc8 | 6j0m | 19.0 | 3.3 | 15 |
| | | T6SS | VgrG1 | 4uhv | 18.7 | 3.2 | 11 |
| | | T4 | Baseplate structural protein Gp27 | 2z6b | 18.4 | 3.2 | 10 |
| | | P2 | | 2wzp | 10.7 | 4.1 | 11 |
| | | RcGTA | | 6teh | 9.2 | 3.5 | 8 |
